# Refined biomimetic model of chronic pain is healed with erythropoietin

**DOI:** 10.1101/051979

**Authors:** Mary R. Hannaman, Douglas A. Fitts, Rose M. Doss, David E. Weinstein, Joseph L. Bryant

## Abstract

Humans suffering with chronic pain may have no evidence of a lesion or disease. They are managed with a morass of drugs and invasive procedures. In many, their persistent pain occurs after the healing of a soft tissue injury, with a neural source hypothesized. Opiates, commonly used to mitigate their symptoms, can cause an increase in neuropathic pain over time. Current animal models of neuropathic pain commonly create direct neural damage with open surgeries using ligatures, neurectomies, chemicals or other forms of intentional trauma. However, we have observed clinically that after an injury in humans, the naturally occurring process of tissue repair can cause chronic neural pain. We show here how the refined biomimetic NeuroDigm GELTM Model, in the mature male rat, gradually induces neuropathic pain behavior with a nonsurgical percutaneous injection of tissue-derived hydrogel in the tunnel of the distal tibial nerve. This perineural model creates a mononeuritis with the biogenic matrix induction of tissue remodeling, the last stage of tissue repair. Repeated behavioral analgesic testing over 5 months in the model implied a unique predictive validity for all analgesics tested. Morphine, initially effective, had an increase in pain behavior over time, suggesting an opioid-induced hyperalgesia, as seen in humans. Celecoxib produced no analgesia, while gabapentin and duloxetine at low doses had profound analgesia. Histology reveals focal neural remodeling, with neural regeneration, as in human biopsies. For the first time, targeted erythropoietin appears to heal neural pain, by extinguishing bilateral pain behavior present for over 4 months.

*translational model, neuropathic pain, erythropoietin, neural regeneration, soft tissue injuries, neuritis, tissue repair, hydrogel, animal model of disease, neural remodeling, age, opioid-induced hyperalgesia, morphine resistance, analgesics, refined pain model, matrix remodeling, neuroinflammation, predictive validity, habituation, estrogen*

## Introduction

The development of chronic neural pain following soft tissue injuries in humans is an uncommon but disabling complication (1–5). The persistent pain usually begins gradually, continuing for months to years, irrespective of the specific cause of the trauma. The typical initiating causes of the antecedent soft tissue injuries include blunt trauma, strains, surgery, industrial injuries, radiation, fractures, vibration, and repetitive motion (6–16). The resultant disuse also contributes to the tissue matrix stiffness, edema and pain (17). Despite a common history of trauma, a trauma-specific neural lesion or occult nerve injury are seldom recognized. Since some patients have the finding of pain long after these injuries have healed, the existence of neural lesions has been implied in this cohort (18). In the absence of an identified neural lesion, many of these patients have been hypothesized to have a neural “generator” or an “ectopic” site (19–29).

The perceived absence of a specific neural injury site, or recognized neural abnormality, does not mean these patients lack such a lesion; these lesions may be “clinically invisible” and below our current level of detection (30), without using invasive techniques. The persistence of pain behaviors in these individuals argues in support of a local neural activation site. In vivo peripheral nerve imaging techniques (31–37) and diagnostics are presently being developed (38), (39), however, most of these imaging techniques cannot yet detect abnormalities in small branches of the distal peripheral nerves (18), which are the fibers most likely to be effected in soft tissue injuries.

A logical cause for the gradual appearance of chronic pain following soft tissue trauma is the predictable changes that occur during the tissue repair process at the affected site. These changes involve the removal of debris, fibrosis, and the regeneration of damaged tissue, including muscle, nerve, vasculature and extracellular matrix. The remodeling of tissue may result in nerve compression, with delayed onset of pain (40). One such example of the ability of minimal pressure on the nerve causing severe pain, is trigeminal neuralgia, where even micro-compression of the nerve root can cause severe pain (41). The timing of the onset of chronic neuropathic pain parallels tissue morphologic events that occur during healing and tissue remodeling of the affected area (S1, (42), (43). We hypothesized that it is during tissue remodeling that an accumulation of matrix and possibly local edema alter the neural microenvironment (44) and contribute to the compression of vulnerable neural fibers, resulting in focal neural injury. These injuries cause atypical forces to intraneural pathology and then abnormal function (45) of peripheral glia and neurons (46), and the subsequent pain syndromes that are observed in these patients. To test this biophysical hypothesis we have created a model of a discrete focal lesion in the rat rear limb that recreates clinical findings similar to those seen in focal neuritises in humans.

Doubts have been raised about whether or not rodents can represent the human condition in neuropathic pain because few effective analgesics have been discovered using them (1), (47). We consider the social behaviors (48), tissue healing (49), (50) and the similar evoked neural pain behaviors that humans share with rodents as confirmative to the relevance for their use. Two critical factors we used in developing this model were the rodent's correlated human age (51) and the clinical relevance of the biologic pathophysiology embodied by the model (1). Presently, animal models with neuropathic pain behaviors are created using forms of direct intentional nerve trauma or irritation, chemicals, drugs, usually with open surgical procedures (52), (53). The most commonly used of these are the Spinal Nerve Ligation (SNL) model (54), the Chronic Constriction Injury (CCI) model (55), and the Spared Nerve Injury (SNI) model (56). These models use ligations, neurectomies or a combination to create pain with sensory and motor debility. While these open surgical models are useful in mimicking direct nerve trauma, they do not reproduce the pathophysiology of the delayed onset of neural pain without debility, as usually happens in many patients with neuropathic pain.

Our report details the creation of a biomimetic pain model based on clinical observations of injured patients with soft tissue injuries. The NeuroDigm GEL^TMp^ Model uses a known physiological tissue process to create a persistent neuritis, with chronic neurogenic pain behavior. Crucially, this model mimics patients clinically, since it has no clinical evidence of either neural injury or physical debility (18), (57). We also explore the tissue repair properties of the biologic, erythropoietin, for targeted healing of the mononeuritis, in a pilot study.

## Materials and Methods

### Ethical statement

The protocol was approved by the Institutional Animal Care and Use Committee of NeuroDigm Corporation (IACUC permit number 1-2014/15) and was in compliance with the guidelines of the 8^th^ edition of Guide for the Care and Use of Laboratory Animals. All efforts were made to minimize the number of animals used and pain and suffering.

### Experimental animals

37 Sprague Dawley 9.5-month-old male rats (Harlan facility in Houston, Texas) were received after being raised within their normal social groups. Their initial weights ranged from 440 to 660 grams, with a mean of 545 grams. Rats are usually sexually mature at 6 weeks and socially mature at 6 months. In this study, the rats’ approximate human equivalent age range starts at about 25 years and ends at 41 years of age (58), (59). The rats had no prior drug exposure. A total of 37 rats were received with 36 rats enrolled after baseline testing. Three rats were removed from study for complications (S2 Dataset), with 33 finishing study.

### Housing and husbandry

Ventilation and housing were per USDA guidelines. Each rat was housed singly in clear, open cages in the same room. The room and individual cages had ammonia sensors (Pacific Sentry). No other animals or rodents were housed in the facility. The cages were changed every 2 weeks or earlier. Bedding was 0.25 inch corncob pellets. Food was LabDiet 5V5R with low phytoestrogens, continuous access. Light-dark cycle was 12 hours, with lights off from 7 PM to 7 AM, except when screening. Maximum lumens at cage level were 20-40; at time of pain behavior testing the maximum lumens were 85-100. The room had no high frequency interference detected (Batseeker Ultrasonic Bat Detector), other than that related to the rats on weekly and as needed testing. Water used was municipal water. In each cage, enrichments were 1) a non-plasticized polyvinyl chloride tube 4” in diameter by 6” in length (Bisphenol A free) for shelter and 2) bedding at an increased depth of 0.75 to 1 inch when dry, to encourage burrowing. Facility was in north Texas. All pain behavior testing were performed in the same room as the rats were housed.

### Study design

After receiving, the rats were acclimated for 15 days with baseline testing (Fig 1), then randomized and blinded. The rats were housed singly to limit fighting and tactile social interaction. The rats were randomly assigned to one of 3 groups: GEL procedure, sham procedure, or control (no procedure). The investigator performing the procedures and behavioral testing was blinded to the rat group assignment. Another experimenter did the random group assignment, before the initial procedures. The rats were housed in a separate room during the procedures, and handed to the investigator by an assistant. Tail identification was masked prior to performance of the procedures. The investigator did not know the group assignments at any time, until the unblinding on postprocedure day 149. The locations of the animals on the rack were randomly changed every 10-14 days. Anesthesia of 2-3% isoflurane was administered with the procedure for approximately 2 minutes, after induction. Isoflurane gas was used due to brief anesthetic time needed, enabling less recovery time compared to most injectables. The analgesics effects of each of the four agents were screened 3 times during the 5-month study. The screening involved testing the plantar hindpaws of the rats with standard stimuli used to detect neuropathic pain behaviors. Each rat was tested singly on a wire mesh with manual application of stimuli, for all behavior testing. The behavioral testing was performed usually between 1 PM and 11 PM. Animal welfare observations of behavior, coat and movement were checked daily. Monitoring for signs of infection, water and food use were conducted at least three times a week. Weights were monitored every 4 weeks, or every 2 weeks as indicated. The same female investigator performed all procedures and screenings, with no one else in the room.

**Fig 1.**
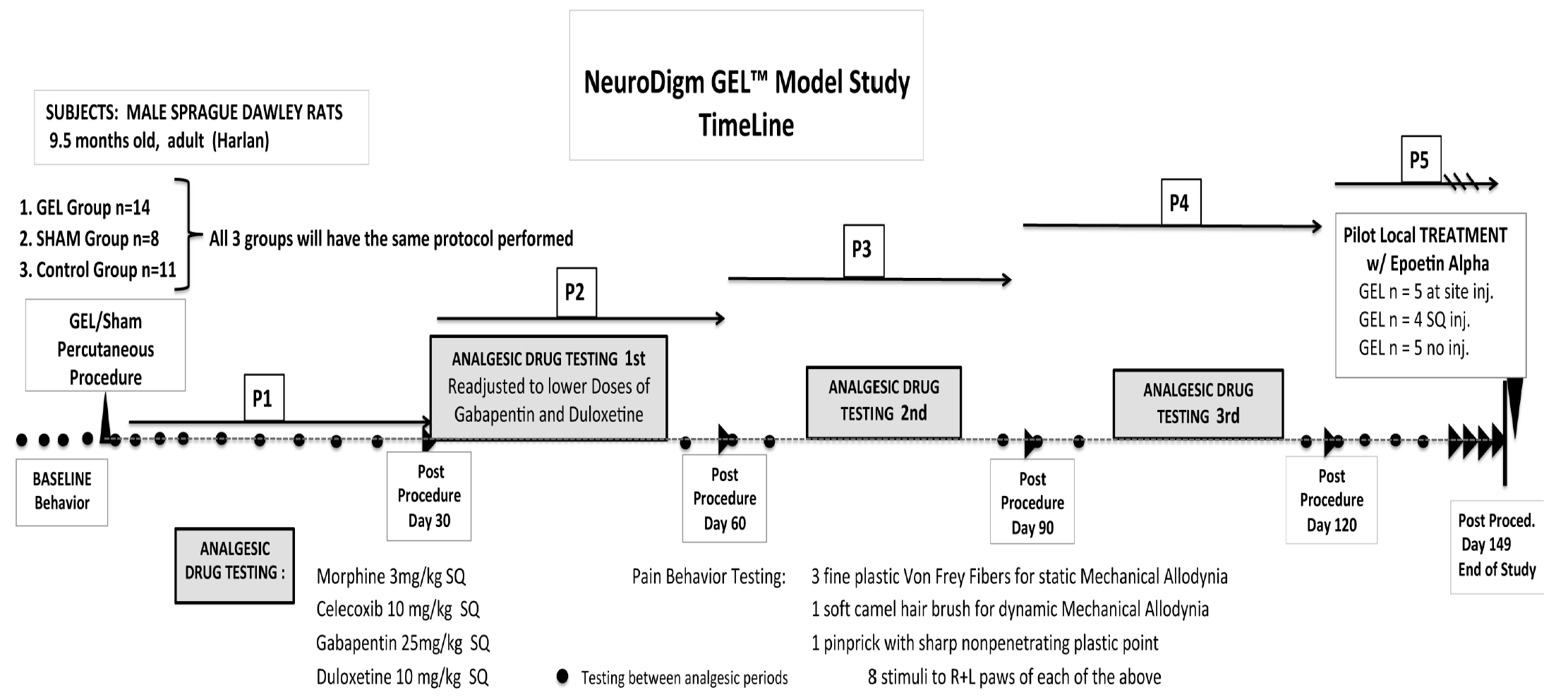
Timeline: Three groups (GEL, sham and control) had four drugs screened for analgesia 3 times over the 5-month study. At the end, a pilot study of a local injection with erythropoietin was performed with the GEL group at the original nerve procedure site.

A blinded pilot study was performed on the effect of a localized application of epoetin alfa (erythropoietin analog, EPO) injected near the GEL^TM^ neural procedure site on post-procedure days 150-160. The four analgesics screened were administered by subcutaneous injections, rather than oral gavage. The method of subcutaneous injection with a custom handheld restraint was less stressful, and had fewer side effects for the rats than oral gavage. ARRIVE guidelines for reporting on animal research were followed and included in the S5 supplement.

### Sample size

The study was designed to test the null hypothesis (60) that a GEL procedure does not differ from the controls during 5 months after procedure on the dependent variables of paw withdrawals in response to von Frey fibers, a camel-hair brush, and pinprick. The alternative hypothesis was that, over time, there is a difference between the groups. The experiment was designed to discover the smallest biologically important effect, optimizing the number of animals used (61). Using data from 3 prior rat experiments (n =55), base sample sizes of 8 per group were selected in order to provide 95% power to detect a difference of 1 paw withdrawal on average between the GEL and control groups with a Type I error rate of .05 if the standard deviation in response to the lightest von Frey fiber was 0.5 paw withdrawals. An additional 3 animals per group were added to compensate for a possible loss of sample size during the 5-month study, and an additional 4 animals were added to the GEL group for aging, illness, technical complications, and a pilot study with local EPO. With the only constraint being the final sample sizes, the animals were otherwise randomly assigned to one of the three groups before the procedures were performed. Final sample sizes: GEL n =14, control n =11, sham n=8.

## Experimental procedures

### Percutaneous procedure for GEL and sham

During isoflurane anesthesia the percutaneous injection procedures were performed. The hydrogel used in the GEL group was the proprietary biological NeuroDigm GEL™, that is composed of purified biocompatible tissue-derived by-products and peptides of mammalian soft tissue, as found in the perineural tissue milieu after a soft tissue injury. Such injectable implant products are used in human surgeries, dermal procedures, and wound healing, with rare reactions of acute inflammation. Purified biological proteins and hydrogels are normally absorbed by the tissue they are implanted in, over days to weeks, and are rarely antigenic (62), (63). The hydrogel we used was introduced within the left (ipsilateral) tibial neural tunnel below the popliteal area at mid lower leg, with aseptic technique. First, the skin was pierced with a sterile 19 gauge needle tip; then a sterile, custom tapered, blunted 21 gauge hollow probe entered the skin puncture site, to gain access to the tibial nerve tunnel. Methods for accessing the tunnels of several nerves are described in U.S. patents 7015371, 7388124. Since the rats were older, with knee contractures, a distal to proximal tunnel access was used in this study. The point of the probe's percutaneous entry was over the Achilles tendon, proceeding proximally subcutaneously, then it pierced the fascia between the distal origins of the medial and lateral gastrocnemius muscle and entered the anatomic tunnel posterior to the tibialis posterior muscle and medial to the soleus, where the tibial nerve courses. Upon entering the neural tunnel, the probe was softly glided in avoiding resistance or nerve contact. In the mid-tibial tunnel of the lower leg 0.3 cc of the GEL^TM^ or the sham’ s' Ringer's lactate was deposited, and then the probe was withdrawn.

### Primary Outcome measure

The outcome measures of light touch and pinprick were chosen since they are used clinically in evaluation of pain in humans. The primary outcome measure for neuropathic pain used in this study was the elicited behavior of paw withdrawal to stimuli for mechanical allodynia and mechanical hyperalgesia. Hypersensitivity to the stimulus of non-noxious light touch or mechanical allodynia was tested with 3 measured forces of von Frey filaments of 2 grams, 6 grams, and 10 grams and a soft fan shaped brush. Exaggerated response to normally painful stimulus, or mechanical hyperalgesia, was tested by pinprick, with a custom calibrated, non-penetrating, sharp plastic tip.

### Procedure to elicit neuropathic pain behavior

For static mechanical allodynia von Frey filaments (Semmes Weinstein Mono-filaments North Coast Medical TouchTest®) exerting forces of 2 grams, 6 grams and 10 grams were used. Dynamic mechanical allodynia was tested with a fan sable brush (09004-1002; Dick Blick Art Materials). Mechanical hyperalgesia was tested with a custom sharp non-penetrating plastic point calibrated to elicit 2-4 paw withdrawals at baseline. On routine behavior testing paw withdrawals to each of the 5 stimuli were recorded. The stimuli were applied in the order of: von Frey 2g, 6g, 10g, brush and lastly pinprick. Each von Frey stimulus was applied to the mid plantar hindpaw for approximately one second until the fiber bent, or the paw was withdrawn. The brush was stroked gently from rear to front plantar hindpaw, and the pinprick stimulus tip was touched to the mid plantar hindpaw until the paw withdrew or the skin visibly indented. Each stimulus lasted about one second. Each stimulus was applied 8 times to the hindpaws bilaterally; contralateral first, then ipsilateral. Time between each stimulus application was usually 2-4 seconds or longer. For each stimulus the total number of hindpaw withdrawals was recorded as a data point.

### Analgesics administration

The analgesics were administered by subcutaneous injection (27 g 1.5”), with a custom administrator-held restraint device to reduce handling and stress. Morphine (West-Ward) was mixed with normal saline and administered at a dose of 3 mg/kg one hour prior to screening the analgesic. The vehicle used in mixing the following three drugs was 0.25% methylcellulose (Methocel^®^ A4M Premium LV USP). These three drugs were mixed 24-48 hours prior to use. Celecoxib (Cayman Chemical) was dispensed at a dose of 10 mg/kg one hour prior to screening; gabapentin (Cayman Chemical) was dispensed at a dose of 25 mg/kg two hours prior to screening; and duloxetine (Cayman Chemical) was mixed (mechanically agitated) and administered at a dose of 10 mg/kg two hours prior to screening. The experimenter knew the drugs being screened; the groups were blinded throughout experiment. Injected volumes were less than 1.2 cc.

The original doses chosen for gabapentin and duloxetine had adverse effects in this study, interfering with the testing of pain behaviors. Gabapentin at 60 mg/kg had marked ataxia in all rats, with their hindpaws not staying on testing screen due to lumbering gait and falls. Duloxetine at 30 mg/kg was noted for marked “frozen” hypoactive posture, with increased tone and alertness (no central sedation) to normal handling and testing. After duloxetine was given at this dose, paw withdrawals were not elicited with any stimuli in any of the 3 groups, including the controls. Due to these adverse effects, lower doses were tested and used, as described above. These lower doses had no observed drug side effects and improved the ability to test paw withdrawals (64).

### Erythropoietin treatment injection

In the pilot study, an erythropoietin analog, epoetin alfa (Amgen), was locally administered under isoflurane anesthesia as described for GEL and Sham procedures. The EPO was diluted with normal saline, and 200 units (0.3 cc vol.) was the administered perineural infiltration dose. The dose was administered with a 27 gauge 1” needle near the original GEL procedure site on the left leg (ipsilateral). The original injection approach on the 5 rats at the GEL site was posterior to anterior at mid tibia through the bellies of the gastrocnemius muscle aiming for the tibial nerve tunnel. 2/5 rats had no decrease in paw withdrawals from the original method of EPO injection. These 2 rats had an adapted lateral approach to improve localization of the perineural infiltrate near the GEL procedure site. This adapted injection was through the lateral gastrocnemius muscle targeted to the mid tibial tunnel at lower leg,

### Histology

At the conclusion of the study, three rats were chosen randomly from each of 3 groups: 1.) GEL procedure rats 2.) sham procedure rats of the 5/8 that displayed late onset robust pain behaviour, and 3.) controls. The GEL group had 2 rats that were controls in the EPO pilot study, and one that had received the subcutaneous EPO injection, with no change in pain behaviour noted. The animals were anesthetised, and then perfused with rat Ringers solution, followed by perfusion fixation with 4% paraformaldehyde in Phosphate Buffered Solution (PBS). Following fixation, the lower limb on the operative side was grossly dissected, to reveal the gastrocnemius muscle thus providing a landmark for locating the tibial nerve. Once identified, the distal tibial nerve (below the popliteal area) was dissected free of the surrounding muscle and fascia, and placed into ice-cold 4% paraformaldehyde in PBS for overnight incubation. The following day the paraformaldehyde was replaced with 30% sucrose to cryoprotect the tissue. The cryoprotected samples were embedded in Tissue-Tek OCT (Sakura Finetechnical, Japan) and frozen on dry ice. Cryosections (10¼m) were then prepared and mounted onto SuperFrost/Plus slides (Fisher Scientific, Rockford, IL). Sections were then fixed in 10% Neutral Buffered Formalin for 10 minutes, washed for 5 minutes in 1X PBS to remove OCT, and rinsed with tap H_2_O. Subsequently, sections were then stained in Hematoxylin (Fisher Scientific) for 5 minutes and rinsed with tap H_2_O, differentiated in acid alcohol (1% HCl in 70% EtOH) for 30 seconds and rinsed extensively with tap H_2_O, blued in 0.2% aqueous ammonia, rinsed with tap H_2_O, and stained with eosin (Fisher Scientific) for 1 minute. Sections were then dehydrated by sequential submersion in graded 75%, 95%, 100% EtOH for 5 minutes each, and a final submersion in xylene. The slides were air dried before mounting with Permount (Fisher Scientific) and adding coverslip. Sections were viewed and the images captured on a Nikon 80i microscope, outfitted for digital light micrographs.

## Statistical methods

### Pain behavior statistical analyses

As described in detail in the results section, the data were inspected for compliance with the assumptions of ANOVA. Two areas of concern were noted, particularly the heterogeneous variances in the pinprick data and the very large number of pairwise comparisons that could be compared. The former occurred because (a.) GEL animals that developed pain symptoms tended to score a maximum number of withdrawal responses (8) out of the possible number of stimuli presented (8) leading to some cells with very small or zero variance in the GEL group only, and (b.) the animals in the sham group were not homogeneous in their response to the sham procedure and this greatly increased their variance. We proceeded with the ANOVA for pinprick because of the convenience of describing interaction effects and for comparison with the allodynia data. We note that the pinprick variable was the least likely to generate errors of inference because of the very large effects and consequent minuscule *p* values obtained. Individual sham data are plotted in a separate graph to illustrate the problem there. Type I errors were reduced by testing only planned comparisons among a relatively small number of means and by combining data where appropriate before analysis so that fewer comparisons would be made.

Analyses of the paw withdrawals in response to von Frey fibers, the brush, or the pinprick on the routine test days were conducted using a mixed model ANOVA with one between groups factor (11 controls, 14 GELs, and 8 shams) and two repeated measures factors. The first repeated measures factor was the time the data were collected, with an average baseline period of 4 days prior to the procedures and the five 30-day periods following the procedure, see Fig 1). In graphs these are referred to as post-procedure Monthly Periods 1 through 5 (P1, P2, P3, P4, P5). For analyses, the data point for each animal in each monthly period was the mean of at least 4 routine pain behavior testing days during that month. The second repeated measures factor for the von Frey fiber analysis was a composite factor combining the 3 levels of fiber force (V1, V2, V3) and the 2 sides for a total of 6 levels of different forces tested on bilateral hindpaws. Differences owing to fiber force and sidedness were determined by comparing means with planned comparisons. The second repeated measures factor for the brush and pinprick were the bilateral hindpaws. In the global ANOVA, a *p*value of <.05 was considered significant. Except as noted for the von Frey analysis, planned comparisons were conducted using Fisher's Least Significant Difference test after a global ANOVA was determined to be significant at the .05 level with a two-tailed test. The S2 Dataset Supplement was used in all studies.

### Analgesic statistical analyses

Experiments were conducted with four analgesic drugs administered shortly before the usual testing with von Frey fibers, the brush, and the pinprick. The four drugs were each tested 3 times during the post-procedure period from day 28 to day 149. The effects of the analgesic drugs were analyzed on two dependent variables instead of five (allodynia measures summed together as one variable and the hyperalgesia measure of pinprick as the other). The data were analyzed using a mixed model ANOVA with the three experimental groups as a between-subjects factor and side (left or right) and days (three pairs of pre-drug and analgesic drug days) as repeated measures factors. The effect of the analgesic drug for each pair of days was analyzed using planned comparisons. These comparisons used Fisher's Protected Least Significant Difference test if the corresponding F-ratio was significant or using a Bonferroni-protected contrast if the F-ratio was not significant. All tests used a two-tailed significance level of .05. In all studies the S2 Dataset Supplement was used.

## Results

### Behavioral observations

The rats undergoing the GEL^TM^ and sham Procedures with the isoflurane anesthesia recovered in their cages. They were visually monitored for behavior one hour after procedure. All rats had recovered from anesthesia within 5 minutes and were walking normally without altered gait. Following recovery from anesthesia the subjects did not demonstrate observable pain behaviors (65) or clinical evidence of tissue injury. Throughout the duration of the study all the rats were observed to have normal gait and were without visible evidence of inflammation, swelling, weakness, deformities or positional changes noted on the operated hindfoot, at any time. There was no observed evidence of acute nocioceptive pain, even after the procedures. Their grooming activities were normal. Among the GEL rats 14/14 rats had markedly increased paw withdrawals to pinprick, von Frey fibers and brush by day 23 post-procedure; the paw withdrawals to pinprick became more exaggerated over the remaining months. By post-procedure day 60, 5/8 of the sham rats had developed marked paw withdrawals. The most common reaction to stimuli was a reflexive flinch with most paw withdrawals. Other reactions appearing one month after increased pain behaviors included prolonged shaking and or licking of their affected ipsilateral paw. Similar patterns of paw withdrawal reactions occurred on testing the contralateral side as pain behavior appeared. No chewing of the paws occurred. Behavioral screening for allodynia and hyperalgesia started on postprocedure day 2 and continued until day 149. Each drug was screened 3 times during this period for its analgesic effect on response to stimuli (Fig 1). Results are presented below with the statistics; also see S3 ANOVA Supplement for a comprehensive description of the basic pain behavior statistical results without drugs or EPO.

## Results of behavioral data

### Pain behavior analyses

Before attempting a statistical analysis, we plotted the raw routine days data (without analgesics) of each group for inspection. The very small effects observed with the allodynia data (each of 3 von Frey fibers and the brush) indicated that an individual routine days mean in the GEL group could not reliably be expected to differ from the control group. In order to observe more reliable differences, the data could be combined in one or both of two ways. First, the scores for all four allodynia measurements could be summed together to make a summary variable (i.e., summing data for all 3 von Frey fibers and the brush into a single number representing allodynia). Second, different days of testing for each stimulus could be summed together for an individual variable (such as data for each individual von Frey fiber summed together over 4 testing days to make a monthly average). These methods have different advantages. Plotting each day's data is useful for determining the precise timing of when effects emerge during the long period of testing. Plotting monthly data is useful for observing the small effects that are apparent between the individual von Frey fibers. Information about effect sizes will be presented for the routine days data of the summed variable (von Frey plus brush), and formal statistical analysis will be applied to the much smaller number of means in the monthly data for each individual variable.

### Days of data pattern of allodynia and hyperalgesia after GEL procedure

The data for all routine testing days (no analgesics given) for the combined allodynia variable (von Frey plus brush, top) and for the pinprick (bottom) are presented in Fig 2) below. The most obvious effect is the increase in pain behaviors in response to the pinprick, a measure of hyperalgesia, in the GEL group on both the ipsilateral and contralateral sides. The pinprick hyperalgesia effect occurred in every rat subjected to the GEL procedure. Although the pinprick responses required about 23 days to develop, the symptoms, and therefore the opportunity to study those symptoms, persisted for months and showed no sign of waning by the end of the experiment. Under conditions of a null effect, each of two groups would be expected to have a greater mean than the other about 50% of the time. However, the last day on which the control group had an absolutely greater mean was for pinprick post-procedure day 5 on the ipsilateral side and day 23 on the contralateral side.

**Fig 2.**
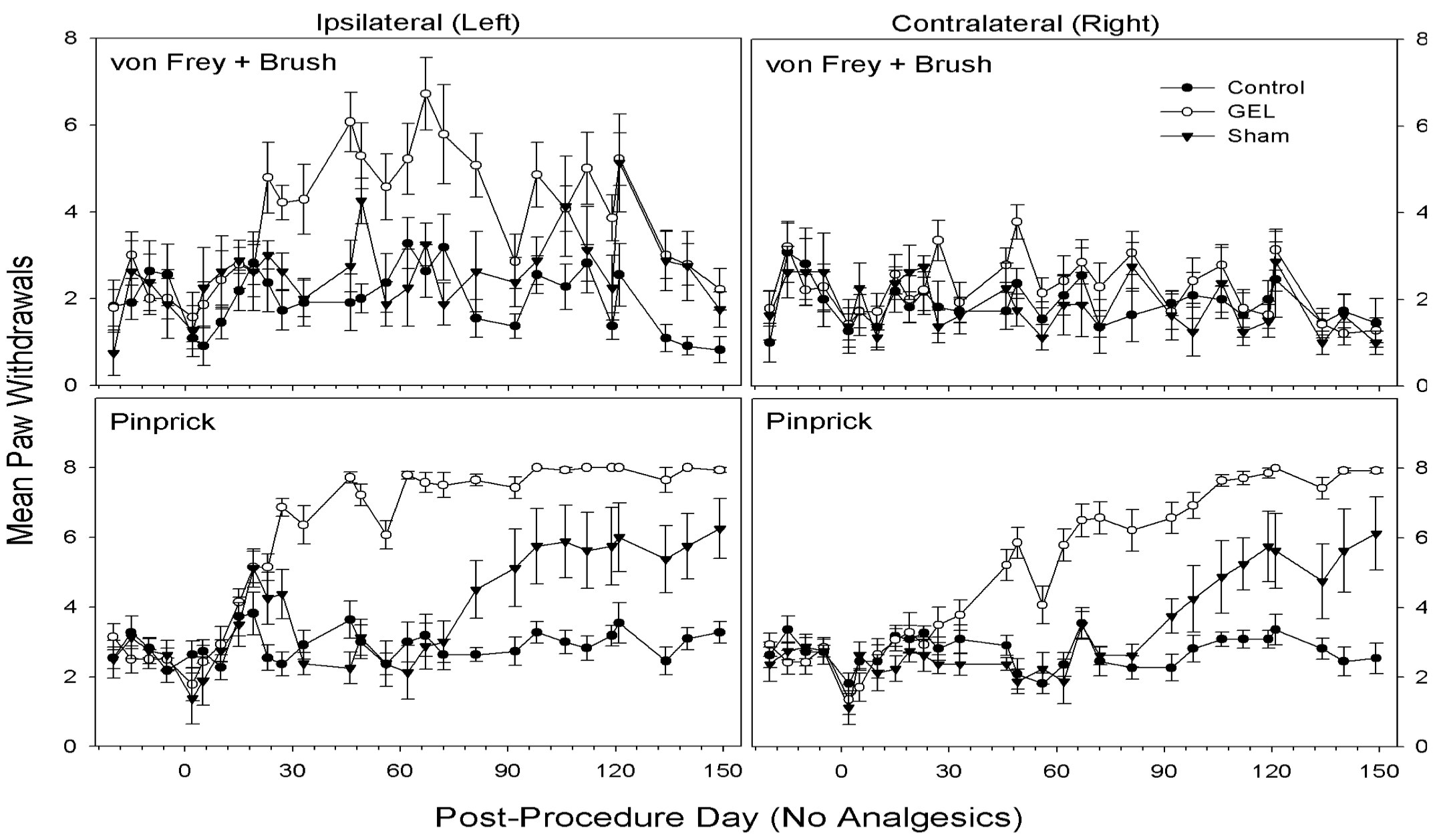
All days of data without analgesics, with the sum of all paw withdrawal responses to the three von Frey fibers and the brush (top) and for the responses to the pinprick (bottom). *The summed allodynia measures on the ipsilateral side were more consistently different on a routine test day basis than any of the individual allodynia measures. Allodynia effects on the contralateral side are not obvious with this type of presentation. Effects related to pinprick hyperalgesia were more robust and persistent than those for the allodynia measures. Shams are not homogenous: 5/8 with pinprick hyperalgesia, 3/8 similar to controls.*

Table 1) provides estimates of effect sizes for the GEL^TM^ effect, which tends to increase with time. The standardized effect size was calculated as the difference between the means of the GEL and control groups on that day divided by the pooled standard deviation of the two groups. These can be used to plan future experiments depending on which interval after the procedure will be studied. Minimum sample sizes are provided to yield at least 80% power in a two-group, two-sided *t*-test with a Type I error rate of .05. Larger sample sizes are required to study allodynia than to study hyperalgesia. More complex designs, such as this one, which include many repeated measurements and multiple groups, will have more error in degrees of freedom for the comparisons than a simple t-test, and will not require such large sample sizes for the allodynia measures.

**Table I.** Effect size (ES) data for hyperalgesia and allodynia on a routine days basis on all routine test control days for the control (C) and GEL (G) groups.

| **Effect** | **<u>Ipsilateral</u>** | **Contralateral** | **Calculated *n*** |
| --- | --- | --- | --- |
| ***Hyperalgesia (Pinprick)*** |  |  |  |
| *Last day C > G* | 5 | 23 | NA |
| *ES > 1.94 SDs* | *day 23-149* | *day 72-149* | <u>6</u> |
| *ES > 2.83 SDs* | *day 46-149* | *day 92-149* | 4 |
| *ES > 3.50 SDs* | *day 72-149* | *day 106-149* | 3 |
| ***Allodynia (von Frey plus brush)*** |  |  |  |
| *Last day C > G* | 19 | 149 | NA |
| *ES > 0.71 SDs* | *day 23-149* | -- | 32 |
| *ES > 0.86 SDs* | *day 112-149* | -- | 22 |
| *ES > 0.91 SDs* | *day 119-149* | -- | 20 |

*Included are the last day that the Control group mean exceeded the GEL group mean, and the inclusive days that the size of effect for the GEL group exceeded the control group by 1.94, 2.83, or 3.50 standard deviations (SDs). ES is the difference between the group means divided by the pooled standard deviation of the two groups. The calculated n is the sample size required to detect a difference between GEL and control groups of the given size for the hyperalgesia or allodynia variable on a routine test day basis in an independent-samples t-test with 80% power and a two-sided alpha of .05. Allodynia was not consistent on the contralateral side. See routine test day means in Fig 2). NA, not applicable.*

Using these data, analgesic screening can start on day 23 or later with a group of n=6 rats using pinprick (mechanical hyperalgesia) on the ipsilateral side. The effect size for hyperalgesia on the ipsilateral side was persistent at more than 1.94 standard deviations from day 23 until the end of the main experiment on day 149. Smaller effects of the GEL group were observed for the combined allodynia variable (von Frey plus brush) than for hyperalgesia with the pinprick, but, unlike the data for the individual von Frey and brush variables, the combined variable showed clear and persistent differences between the GEL and control groups on the ipsilateral side for each testing day. This is important for our subsequent experiments with analgesics. The effect of the GEL procedure is not constant across time, therefore, when testing the effect of an analgesic on a single day with a control value from the same animal, the response must be compared to a control day very close in time to the day the analgesic is given rather than to the average of all control days. To do this successfully, each control day must show a positive effect of the GEL procedure, and this was not true on every day for the individual von Frey fibers or the brush alone. Consequently, we opted to use the combined von Frey plus brush data for all comparisons between individual analgesic and control days (see the section, *Results: Experiments with analgesic drugs;* where statistical analysis of those days are provided).

In order to provide a formal statistical analysis of the individual allodynia variables without summing them together, we instead averaged all days (at least 4) within each month that the animals were tested without analgesics to remove some of the test day variability, stabilize the means, and estimate effect sizes. These analyses are presented in the following sections.

### Mechanical allodynia: von Frey analysis

The data are given in Fig 3) for the three different fiber forces in each group, over all periods applied to both the ipsilateral and contralateral sides. The highest-order interaction of the ANOVA was significant (*F* (50, 750)=2.21, *p*<.001). The control group never significantly exceeded the baseline value in any monthly period on either side. *Asterisks in Fig 3) mark means of the GEL group that were significantly different from both the GEL group baseline and the mean of the control group during the same period.

**Fig 3.**
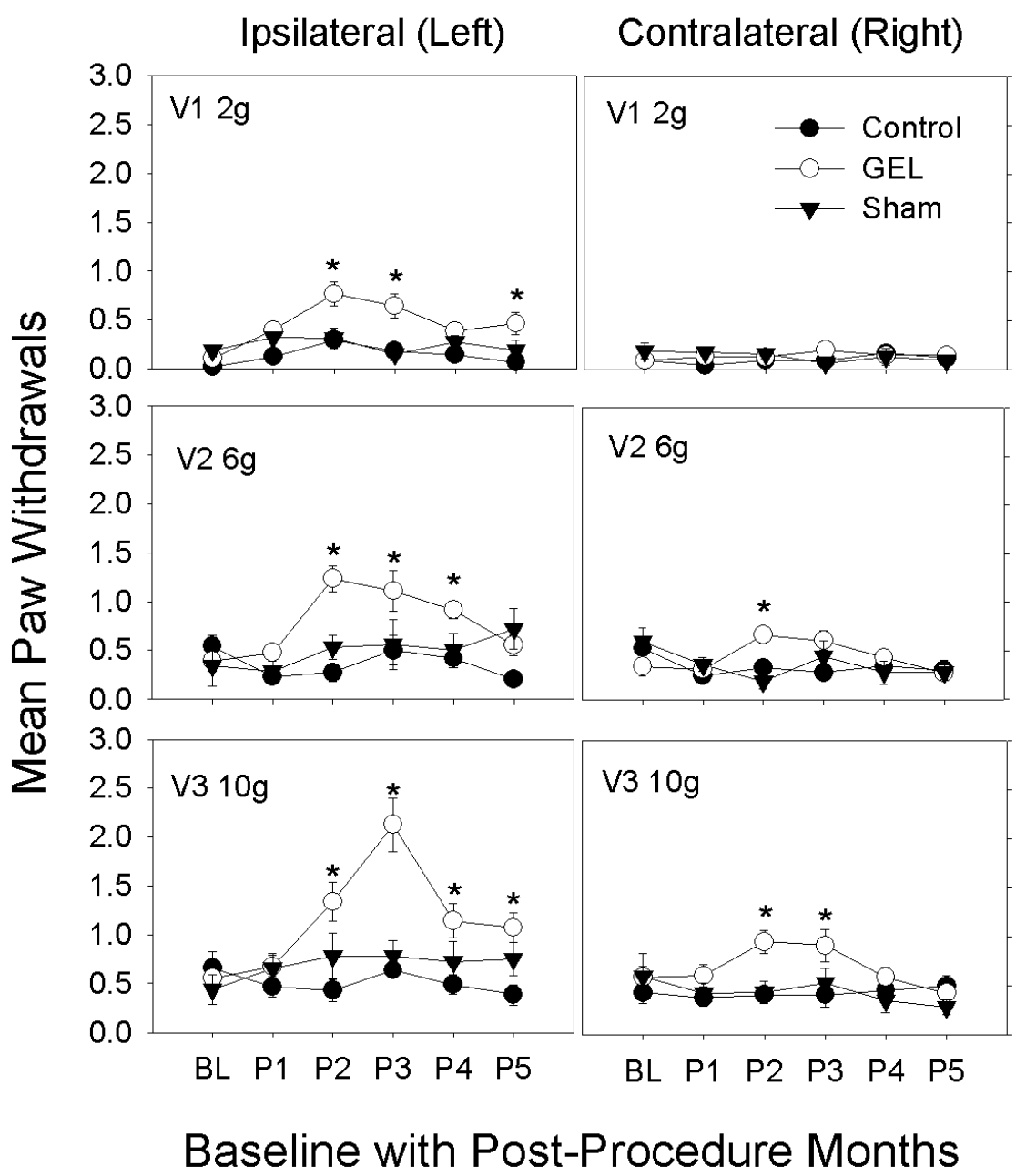
Increased pain behavior of allodynia to light touch with von Frey fibers only seen in the GEL rats, bilaterally; on the left with 2g, 6g and 10g fibers, and on the right with 6g and 10g. This allodynia decreased after the 3^rd^ month for all fibers with this effect. *These graphs depict von Frey fiber results for paw withdrawals ipsilateral and contralateral, for all groups, for three fiber forces over 5 monthly post-procedure periods (P1-P5). Mean and S.E.M. *p < .05 GEL group greater than both GEL group baseline and control group for same period. Reduced responding in the GEL group during period 5 may reflect habituation*.

### Mechanical allodynia: Brush analysis

The GEL^TM^ group had the more robust prolonged pain behavior, of dynamic mechanical allodynia to brush stimuli with increased paw withdrawals, only on the ipsilateral side. This pain behavior peaked by the 3^rd^ month then waned, returning to near baseline by the 5^th^ month. The shams had similar pain behavior that plateaued by the 4^th^ month, persisting until the 5^th^ month at end of study. The response of the shams, with the onset of the pain behavior of mechanical allodynia during the 3^rd^ month that persisted, was not anticipated (Fig 4). The data for the ipsilateral and contralateral paws are presented in the upper part of Fig 4). The highest-order interaction was significant (*F* (10, 150)=1.943, *p*=.044). The control group never significantly exceeded the baseline value in any monthly period on either side. *Asterisks in Fig 4) mark means of the GEL group that were significantly different from both the GEL group baseline and the mean of the control group during the same period.

**Fig 4.**
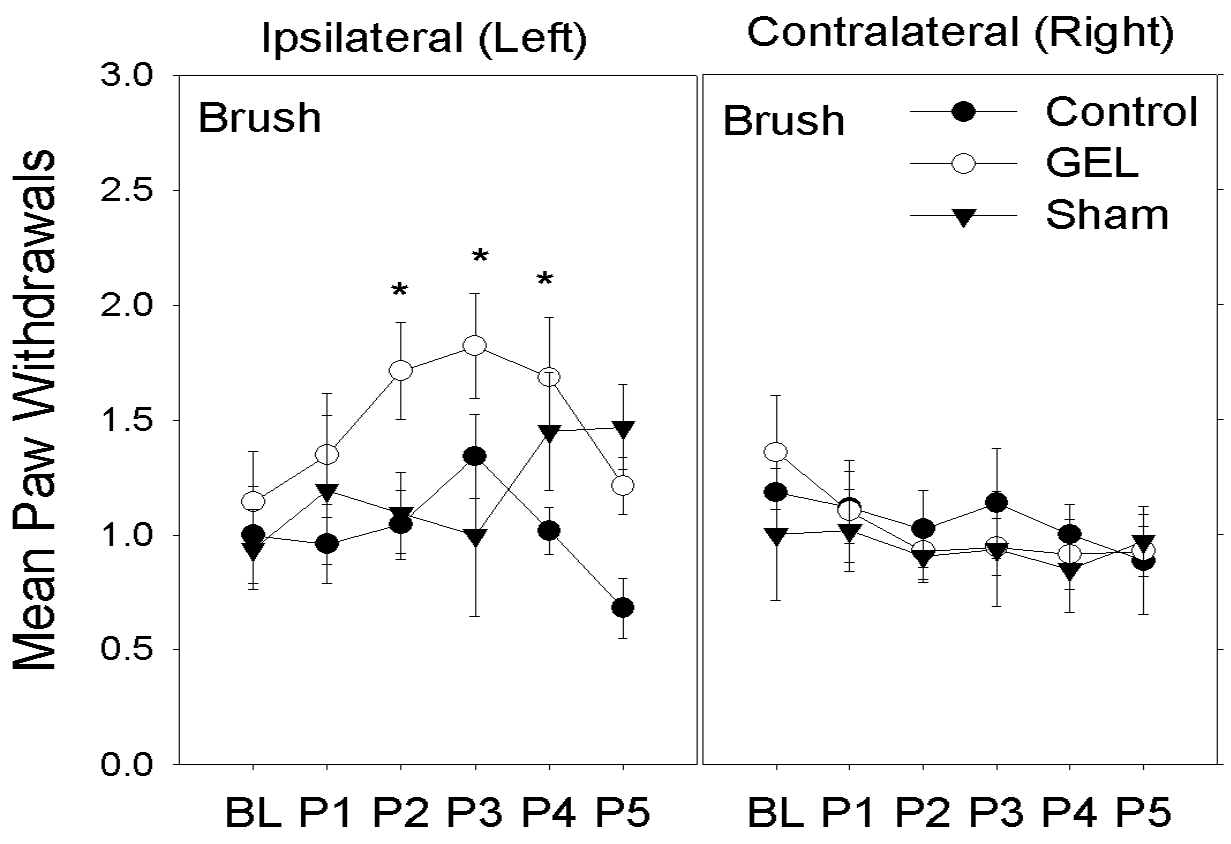
The GEL and sham groups had dynamic allodynia or hypersensitivity to the brush only on the ipsilateral hindpaw. *These graphs depict brush results for paw withdrawals ipsilateral and contralateral to procedure over 5 monthly post procedure periods (P1-P5). Mean and S.E.M. *p <.05 GEL group greater than both GEL group baseline and control group for same period. Reduced responding in GEL and control groups during period 5 may reflect habituation, that is not present in shams, as their allodynia increased in P4-P5*.

### Mechanical allodynia: Combined von Frey and brush analysis

With the combined light touch stimuli, the GEL group had the more robust prolonged pain behavior, of all mechanical allodynia with increased paw withdrawals, bilaterally. This pain behavior peaked by the 3^rd^ month then waned returning to near baseline by the 5^th^ month. The shams had late onset of allodynia pain behavior on the left ipsilateral side that peaked by the 4^th^ month and persisted until the 5^th^ month at end of study.

Fig 5) provides an overview of all allodynia measures by presenting the sum of the paw withdrawals for all three von Frey fibers and the brush combined. Effects are more pronounced in the summary variable than in the individual variables. In this study, the effect sizes owing to the GEL procedure on a monthly basis were .88 standard deviations above the control group in month one and at least 1.5 standard deviations thereafter. Hence, multiple measures summed together and then averaged over several days stabilized the means, reduced the variability, and increased the standardized effect size compared with the day-to-day data in Fig 2) and Table 1). A tendency to a decrease in responding in the last 2 months is apparent in both the control and GEL groups, although the effect between those groups was maintained at 1.9 standard deviations in post-procedure period 5. By contrast, the sham group increased responding during months 4 and 5.

**Fig 5.**
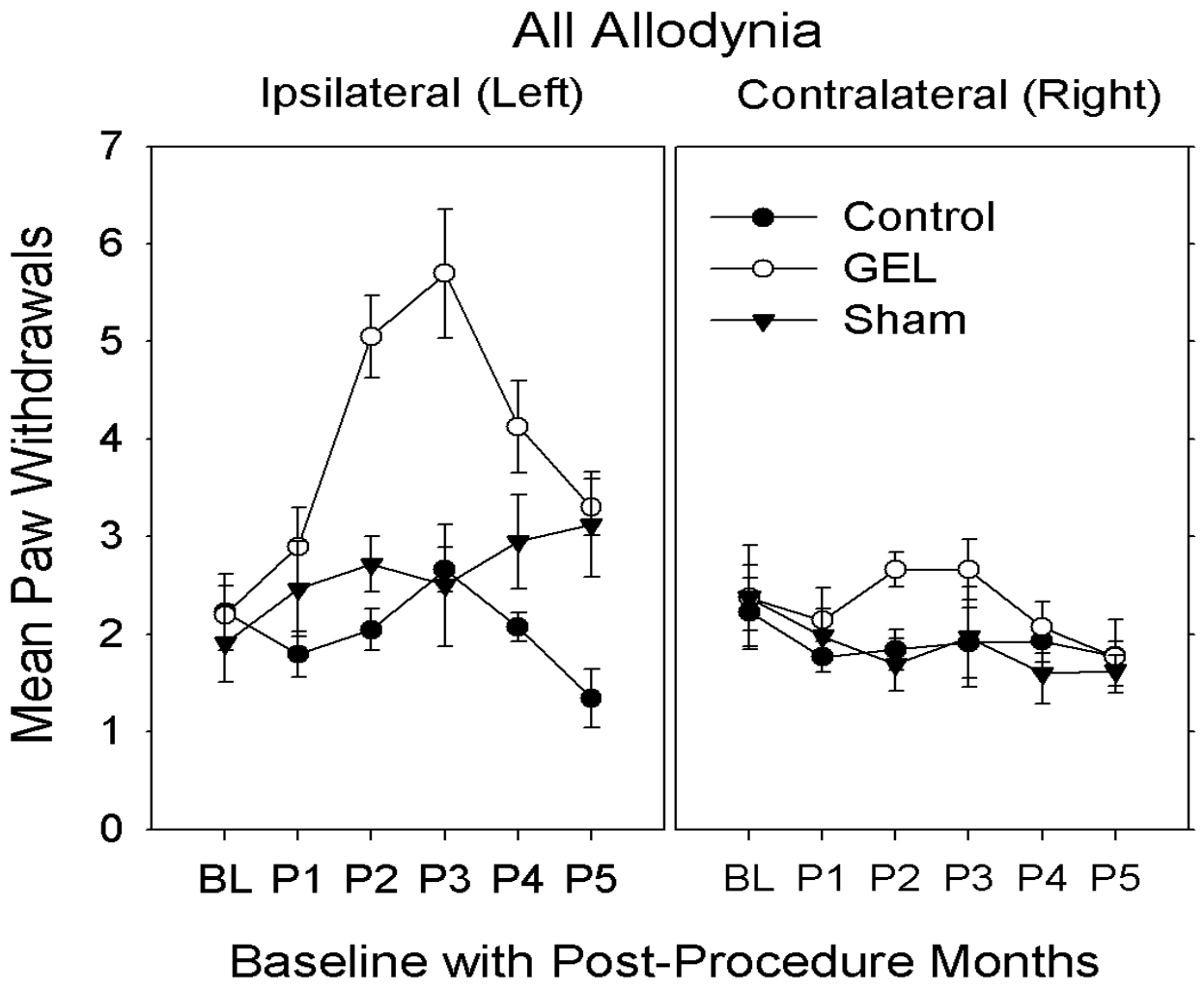
Mechanical allodynia, to the combined (all) light touch stimuli of von Frey and brush, is predominantly in the GEL group, bilaterally, left more pronounced than the right. The shams begin to develop allodynia by the 4^th^ month on the left procedure side. *Reduced responding in GEL and control groups during period 5 may reflect habituation, that is not present in shams*.

### Mechanical hyperalgesia: Pinprick analysis

The GEL group had the earliest and most persistent pain behavior of mechanical hyperalgesia with increased paw withdrawals to pinprick, bilaterally. The hyperalgesia was first present on the left side and within a few weeks present on the right side. This pain behavior was vigorous after the first month and persisted robustly for 4 months, until the end of the study. The shams had similar pinprick pain behavior bilaterally that peaked by the 4^th^ month and persisted until the 5^th^ month, at end of study. The controls had no pinprick pain behavior during the study.

The data for pinprick are presented in Fig 6). The highest-order interaction was significant (*F* (10, 150)=4.592, *p* <.001). The control group never deviated from its own baseline value in any postprocedure period on either side. By contrast, the GEL group's paw withdrawal response on the ipsilateral side was significantly greater than baseline during all 5 post-procedure periods, and the GEL group on the contralateral side was significantly greater than baseline during periods 2 through 5. *Asterisks in Fig 6) mark means of the GEL group that were significantly different from both the GEL group baseline and the mean of the control group during the same period.

**Fig 6.**
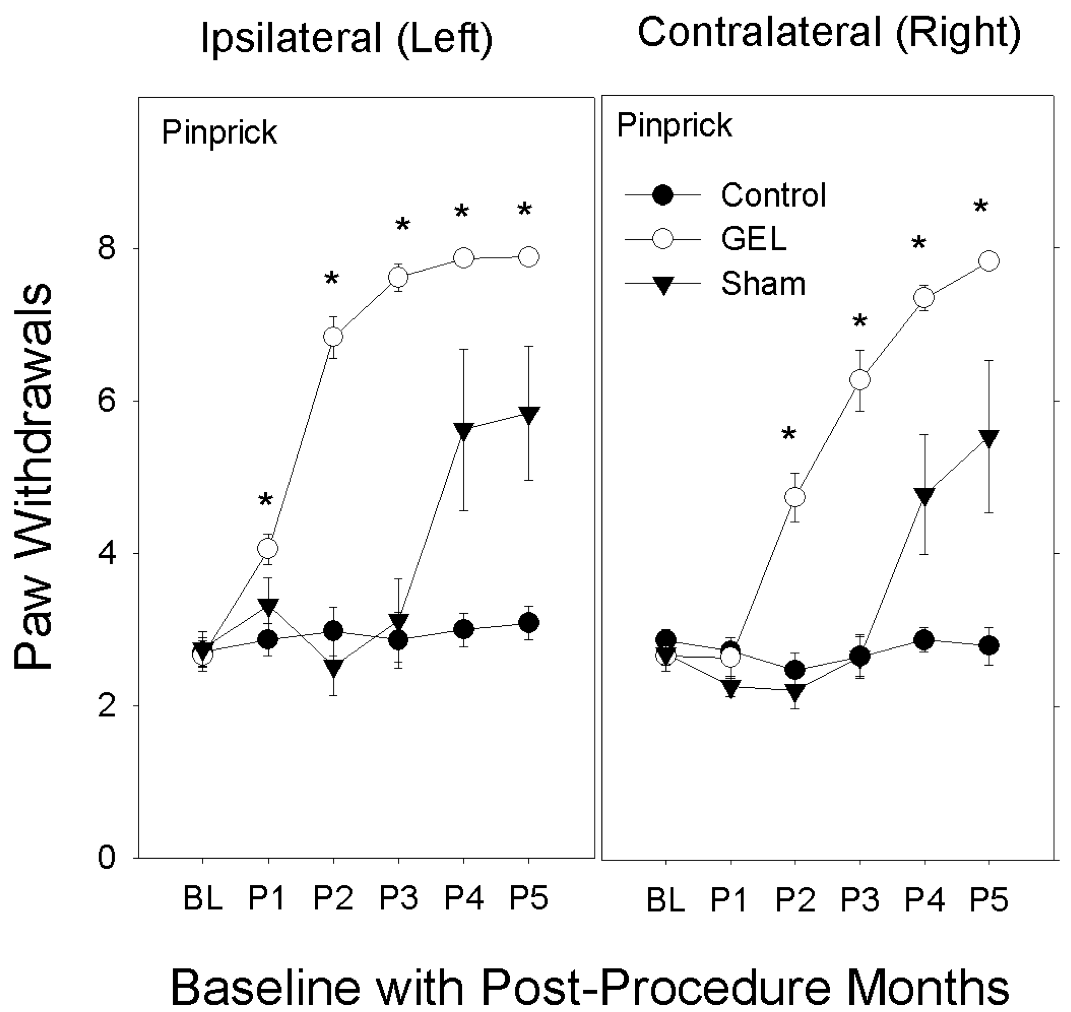
The GEL and later the sham groups develop mechanical hyperalgesia to pinprick bilaterally. 2-3 weeks following the onset of this pain behavior on the left, the same pattern of pain behavior begins to develop on the contralateral side. The graphs depict the sum of all paw responses to pinprick on the ipsilateral and contralateral sides during the 5 monthly post-procedure periods. Shams are not homogenous: 5/8 with pinprick hyperalgesia, 3/8 similar to controls. No habituation effect was noted.

Between-group comparisons for pinprick indicated that the three groups were not significantly different during the baseline period on either the contralateral or ipsilateral side. On the GEL procedure ipsilateral side during the post-procedure periods, the GEL group responses significantly exceeded the means of the control group during all 5 periods. On the contralateral side, the GEL group significantly exceeded the means of the control group during periods 2 through 5.

### Individual data for sham group

Retrospectively, we noted that 5 of 8 sham procedure animals developed pain behavior bilaterally, similar to the GEL^TM^ animals, in post-procedure monthly periods 4 and 5 (after 3 months); and the 3 remaining sham rats behaved similarly to the control group.

The foregoing analysis did not stress any effects that might or might not be different between the GEL and sham groups. The reason for this is that the sham group itself was not homogeneous in the responses of the animals to the sham procedure. Individual data for the sham animals are presented in Fig 7). The data on the shams, in all the pain behavioral studies and in the analgesic screening, included the results from all eight shams.

**Fig 7.**
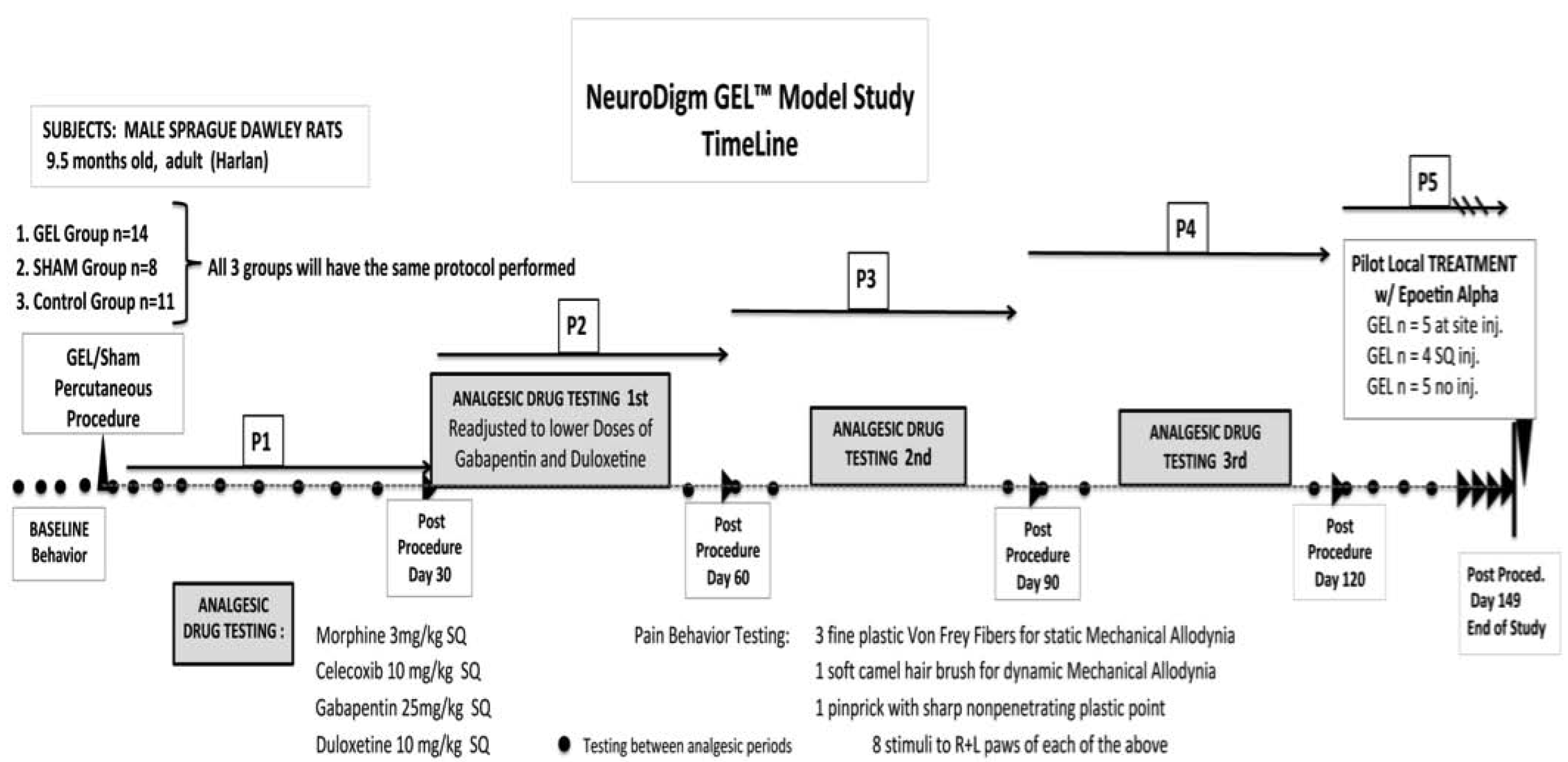
5/8 shams had bilateral pinprick mechanical hyperalgesia during the P3 month persisting until end of study. Pain behavior on right developed 2-3 weeks after the left. *The remaining 3 animals resembled the control group. Graphs depict sum of paw withdrawals for eight individual shams; the sum of all responses to von Frey and brush stimuli (top) or to pinprick (bottom) over baseline and for 5 months. In P4 and P5 for bilateral pinprick there is complete separation between the 3 behaving like controls and the 5 behaving like GEL*.

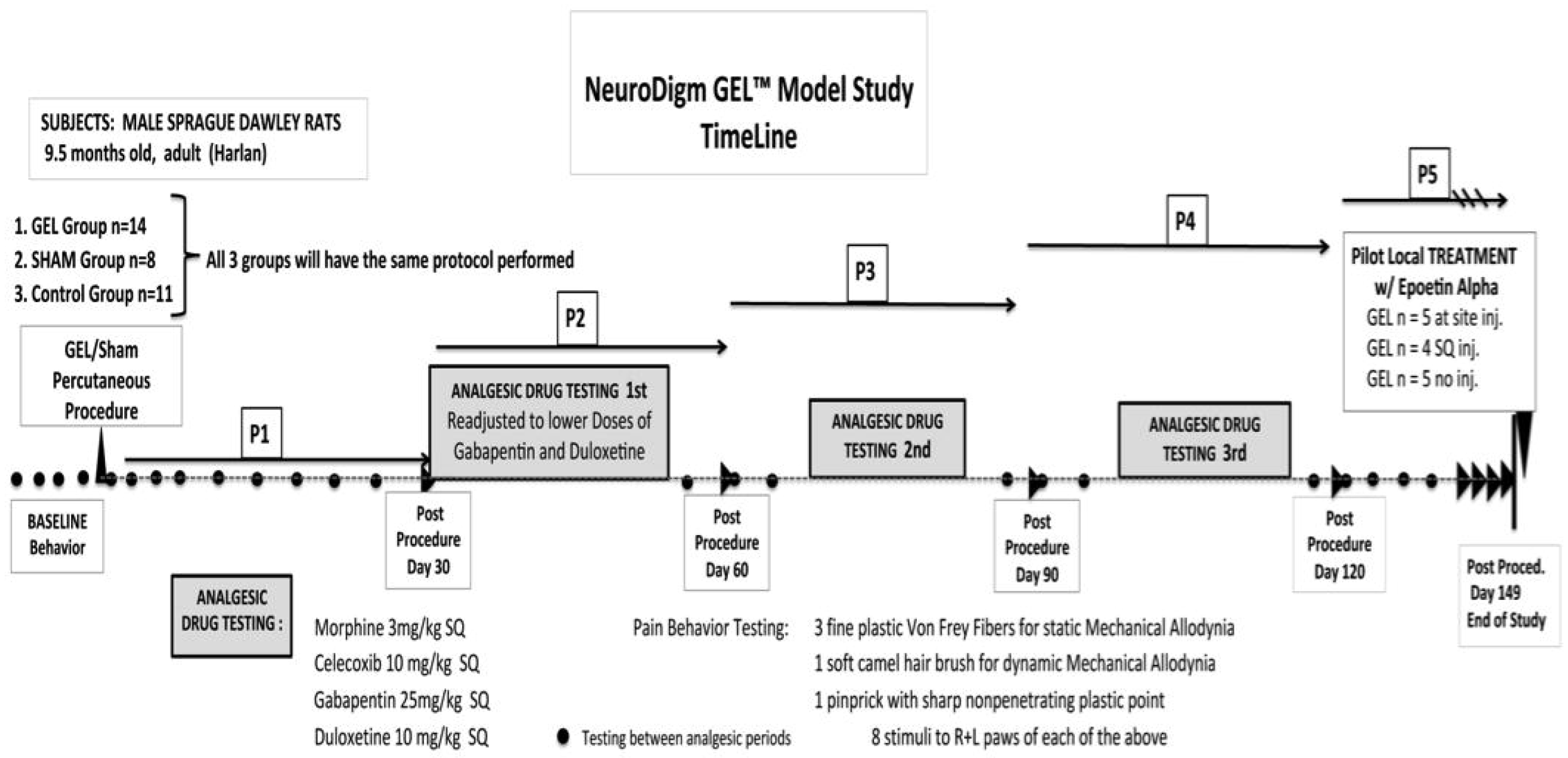

### Summary of pain behaviors

For the GEL procedure group, we found that the gradual pattern of development of ipsilateral pain behaviors was usually followed with the gradual onset of contralateral pain behaviors within 2-3 weeks. In the GEL group, the light touch mechanical allodynia responses had a gradual onset after the first month, decreasing slowly after the third month (P3). Once the pinprick hyperalgesia developed in the GEL group it became robust and persisted until the end of the study. The sham procedure also gradually induced the pain behavior of pinprick similar to the GEL procedure in 5 of 8 rats on the ipsilateral side after day 72, followed by the contralateral side after day 90. The affected shams (5/8) with elicited pain behavior of mechanical hyperalgesia became robust after the 3^rd^ month P3, persisting until the end of the study. The shams did not have a robust response to von Frey or brush stimulation. The control group developed no pain behaviors.

## Results of experiments with analgesic drugs

To control for the effects of time, it was important to compare the data for each analgesic's screening day to a single pre-drug control day, prior to the drug's administration. As illustrated in the top part of Fig 2), an effect on the GEL group's ipsilateral side was apparent on individual control days, when the paw withdrawals for all four allodynia measures were summed together into one variable. For that reason, we analyzed only the composite allodynia variable and the pinprick for responses to analgesics. The data for each analgesic drug are displayed in the figures as bars for the mean response, with the analgesic drug on a particular day (the days listed in the x-axis label) paired with the control data from the GEL group's routine test day one to four days prior. Throughout the following analyses, the mean number of paw withdrawals in the GEL group on the ipsilateral side was significantly greater than that of the control group on all pre-drug days for both the von Frey plus brush variable and for the pinprick. On the contralateral side, the GEL group response was significantly greater than the control group on the pre-drug days corresponding to all analgesic days for the pinprick and on most days for the allodynia variable.

### Morphine

Morphine at 3 mg/kg had an escalating loss of effectiveness bilaterally, which was not due to tolerance as each of the 3 doses were at least 30 days apart. Morphine 3 mg/kg is usually a toxic dose in humans. The data for the morphine test days using the von Frey fibers and brush are presented in the top half of Fig 8). The ANOVA revealed that all main or interaction effects were significant at the .05 level including the three-way interaction (*F* (10, 150)=2.322, *p*=.014). Morphine caused significant decreases in responding compared with pre-drug day on both the ipsilateral and contralateral side only on day 28 as denoted by asterisks in the figures. In three instances, morphine actually increased pain responses, as denoted by red plus symbols. These were the only significant increases in pain behavior caused by an analgesic in the entire dataset for the 4 analgesic drugs. The data for pinprick are presented in the bottom half of Fig 8). The ANOVA revealed that all main and interaction effects were significant including the three-way interaction (*F*(10, 150)=2.655, *p*=.005) and significant decreases are denoted by asterisks. The shams and controls also suggest an escalating pattern of stimuli sensitivity after morphine with pinprick.

**Fig 8.**
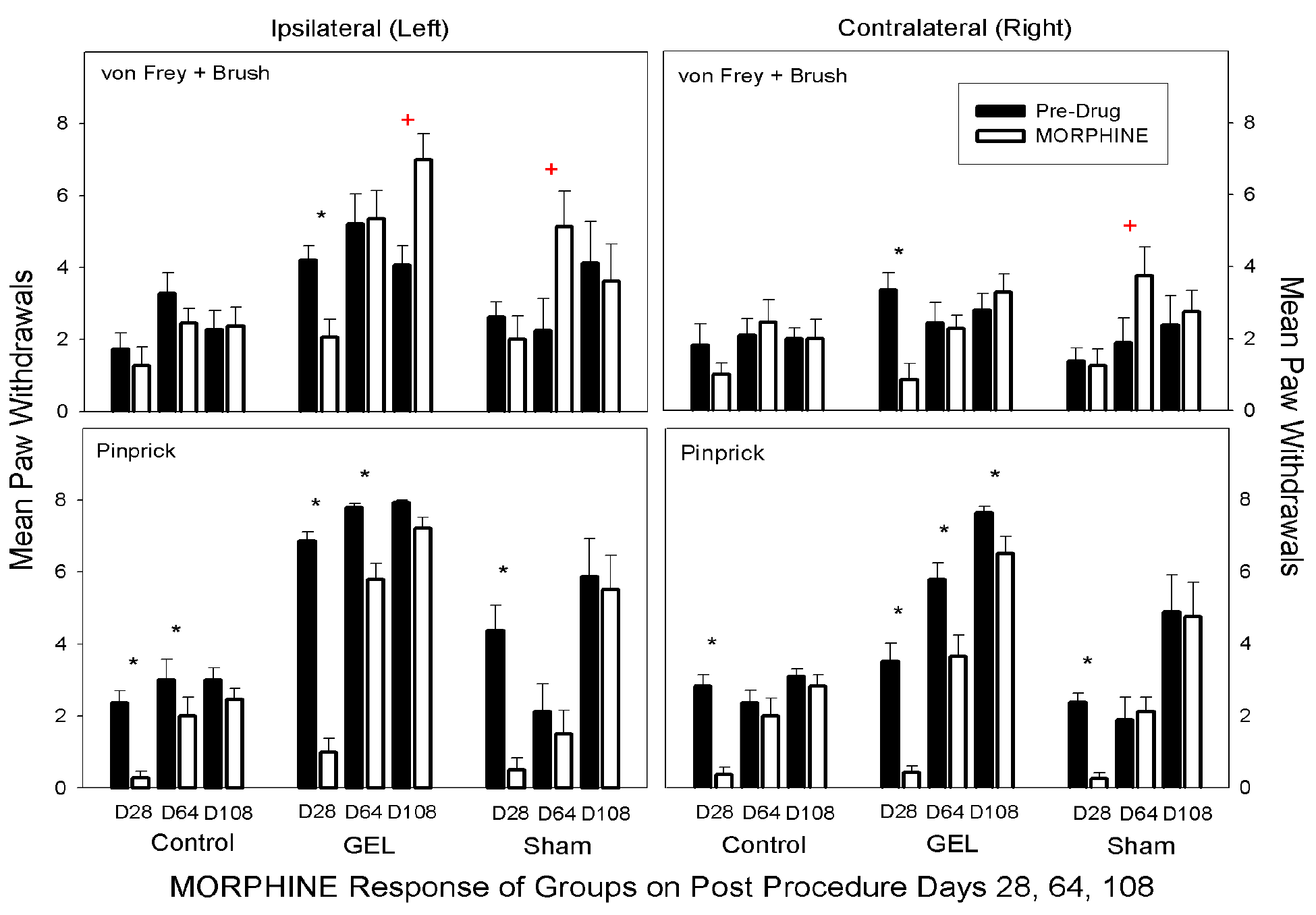
Morphine. Only D28 morphine has marked analgesia with pinprick (bottom) and light touch allodynia (top) across all groups, bilaterally. Morphine became less analgesic in the last 2 testing periods D64 and D108, suggesting a developing opioid-induced hyperalgesia, rather than tolerance. The graph depicts the sum of paw withdrawals on pre-drug control days (black) and on paired morphine dose days (white). Results show the sum of all light touch allodynia measures (top graphs) and the pinprick hyperalgesia (bottom graphs). Mean and S.E.M. *Significant decrease from the paired control day,*p*<0.5. + Significant increase from the paired
control day, *p* <.005.

Morphine was effective early after the GEL^TM^ procedure, but the size of the effect waned with time on both sides. For example, in the GEL group on the ipsilateral side for the allodynia measures, the standardized effect size between the control day and the morphine day changed from a positive analgesic effect of 1.30 pooled standard deviation units on day 28 to no effect on day 64 to a negative effect of −1.25 pooled standard deviation units on day 108. For the pinprick measure, effect sizes were conservatively estimated using the standard deviation for the morphine condition only instead of the pooled standard deviation because of the reduction of the variability as the responses approached a ceiling of 8 paw withdrawals out of 8 pinpricks. The analgesic effect size waned from 4.14 standard deviations on day 28 to 0.64 standard deviations on day 108. A reversal of the effect was not possible for pinprick because the pain response was already maximal on the pre-drug day.

### Celecoxib

There was no analgesic effect of Celecoxib at 10 mg/kg in any group on any day on either side. Celecoxib did not decrease pain behaviors, as demonstrated with a decrease in paw withdrawals. Celecoxib at 10 mg/kg is about 3 times a human dose. Data for the celecoxib days for the von Frey fibers and brush are presented in the top half of Fig 9). The ANOVA revealed significant main effects of groups, side, and drug response and a significant group-by-side interaction (*F* (2, 30)=22.92, *p*<.001). None of the other interaction effects was significant. There was no effect of the celecoxib dose on either side in any group by Bonferroni-protected planned contrasts. The data for pinprick are presented in the bottom half of Fig 9). The ANOVA revealed that all main and interaction effects were significant, including the three-way interaction (*F* (10, 150)=2.675, *p*=.005.

**Fig 9.**
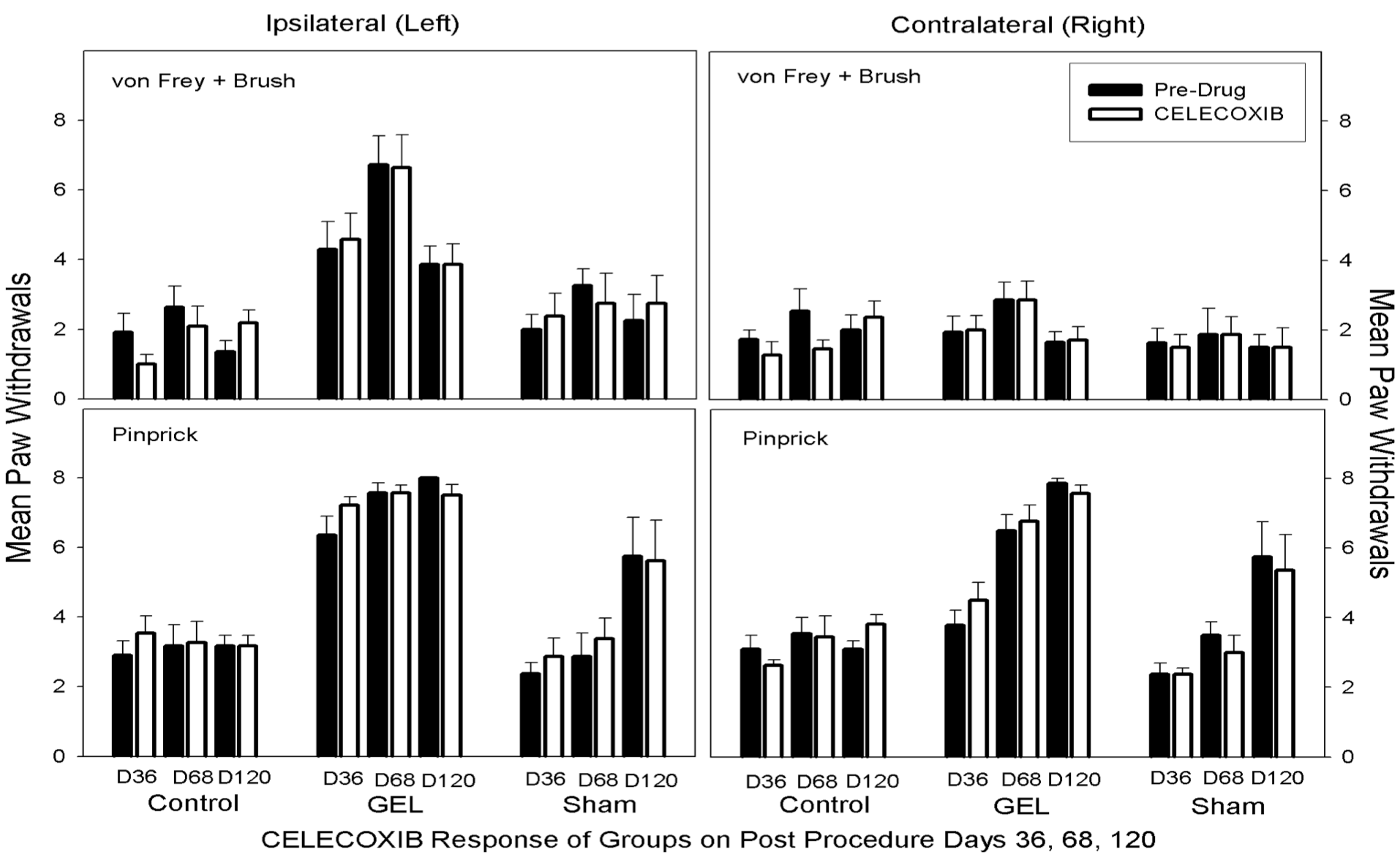
Celecoxib. No analgesic effect of Celecoxib is demonstrated, no significant change in paw withdrawals. Graph depicts sum of paw withdrawals on pre-drug control days (black) and on paired celecoxib dose days (white). Results of behavior testing show the sum of all four light touch allodynia measures (3 von Frey fibers and the brush, top graphs) and the pinprick hyperalgesia (bottom graphs). Mean and S.E.M. Celecoxib did not significantly reduce paw withdrawal responses on any pair of days.

### Gabapentin

On all days on both sides in the GEL group gabapentin at 25 mg/kg robustly reduced paw withdrawal responses in Fig 10). Gabapentin also significantly reduced the level of responding in the control group on days 47 and 76 on both sides. The dose of gabapentin used is nearly equivalent to the human dose. The data for the gabapentin test days using the von Frey fibers and brush are presented in the top half of Fig 10). The ANOVA revealed significant main effects and interaction effects except for the three-way interaction. It appears that the group-by-days pattern of responding was similar on the ipsilateral and contralateral sides, therefore, the groups-by-days interaction term is the important one for analysis (*F* (10, 150)=1.929, *p*=.045). When the data for the ipsilateral and contralateral sides were combined, we found that gabapentin robustly reduced the GEL group responding on all analgesic test days. For comparison to other figures, asterisks in the top half of Fig 10) represent significant decreases from the paired pre-drug day by Bonferroni-protected planned contrasts. By this analysis, the comparison for day 114 on the contralateral side for the allodynia measure in the GEL group was not significant. The data for pinprick are presented in the bottom half of Fig 10). With the exception of the group-by-side interaction (*p*=.055), all main and interaction effects were significant at *p*<.05 including the three-way interaction (*F* (10, 150) =.029, *p*=.03).

**Fig 10.**
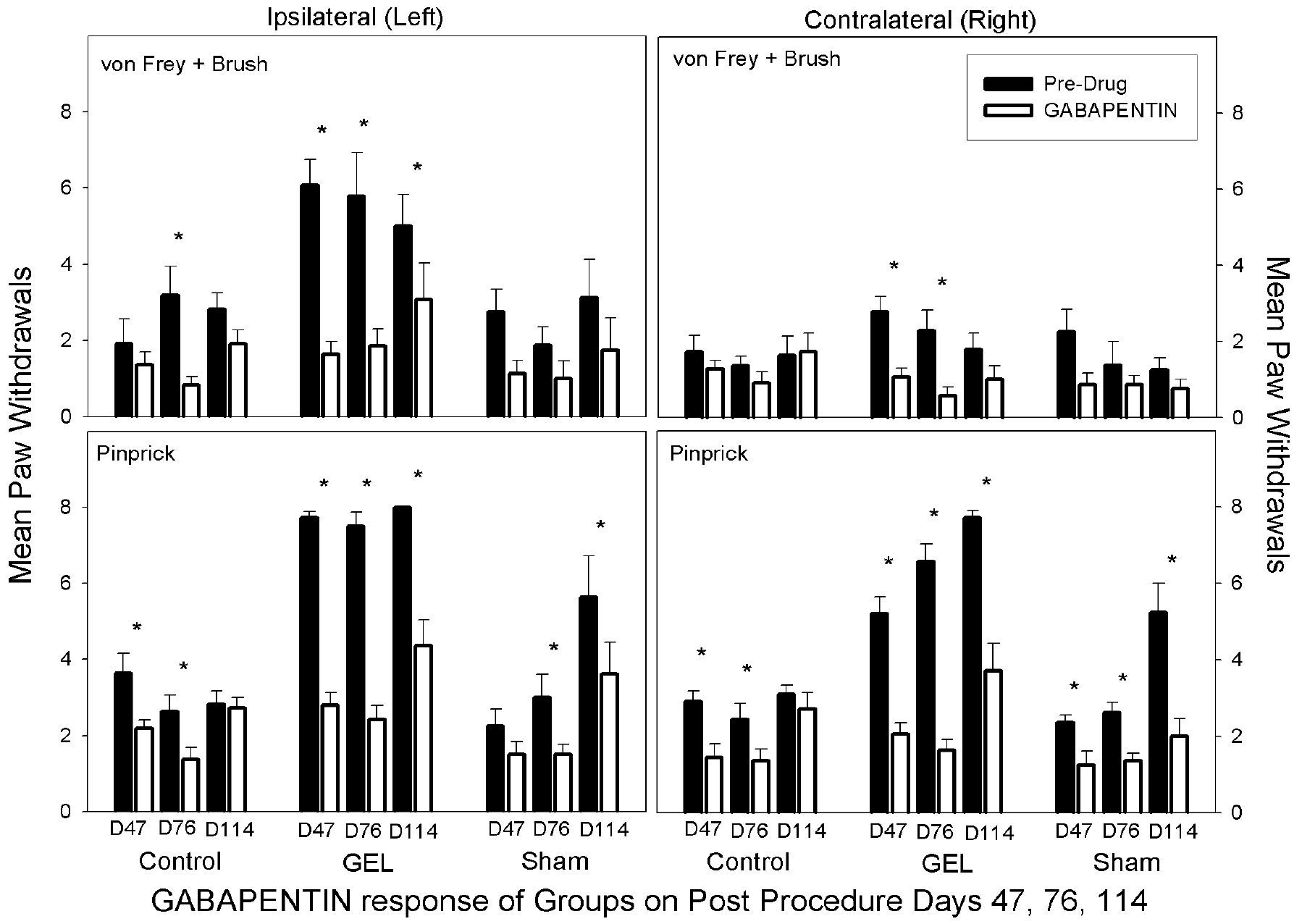
Gabapentin. Analgesia with reduced paw withdrawals on Gabapentin is seen in the GEL group bilaterally with pinprick stimulus and on ipsilateral side with light touch stimuli of von Frey and brush. Graph depicts sum of paw withdrawals on pre-drug control days (black) and on paired gabapentin dose days (white). Results of behavior testing show the sum of all four light touch allodynia measures (3 von Frey fibers and the brush, top graphs) and for the pinprick hyperalgesia (bottom graphs). Mean and S.E.M. 1Significant decrease from the paired control day, p <.05. Gabapentin also suppressed responding in the control group.

### Duloxetine

In the GEL^TM^ group, duloxetine at 10 mg/kg significantly reduced all pain behaviors bilaterally at all time periods. This dose is markedly less than most prior rat doses (64). Duloxetine was very effective as an analgesic at 10 mg/kg; lower more humans equivalent doses (1.5 −2/mg/kg) need studies in this model. The bilateral analgesia was similar in the sham group but exhibited only after the pain behaviors in them began developing after 2 months, and is seen on D83 and D125. Interestingly, duloxetine did not suppress normal responses to pinprick stimuli in the control group as the gabapentin did. The von Frey fibers and brush data for duloxetine are presented in the top half of Fig 11). The ANOVA revealed a significant three-way interaction (*F* (10, 150)=1.99, *p*=.039). The data for pinprick are presented in the bottom half of Fig 11). The ANOVA revealed that all main and interaction effects were significant except for the three-way interaction. The groups-by-days interaction was significant (*F* (10, 150)=11.358, *p*<.001), and the pattern of responding within the groups was similar on the ipsilateral and contralateral sides. Responding in the GEL group was robustly reduced by duloxetine on all days (Bonferroni-protected contrasts).

**Fig 11.**
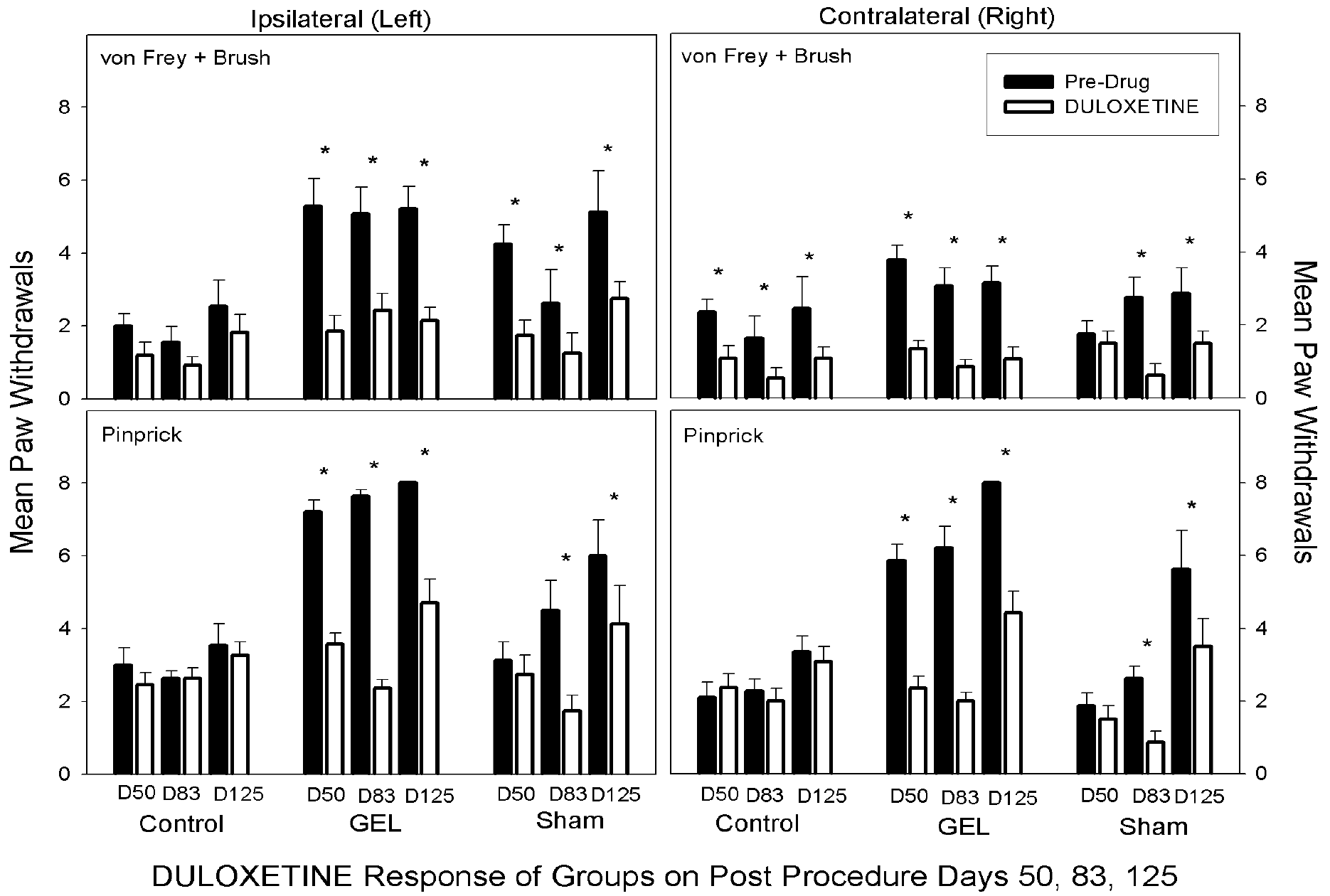
Duloxetine. In the GEL group, duloxetine reduced paw withdrawal responses
consistently throughout the study, demonstrating analgesia to all stimuli, bilaterally. A similar response was noted in the last 2 test days of the shams, D83 and D125. *Graph depicts sum of paw withdrawals on pre-drug control days (black) and on paired Duloxetine dose days (white). Results of behavior testing show the sum of all four allodynia measures (3 von Frey fibers and the brush, top graphs) and for the pinprick hyperalgesia (bottom graphs). Mean and S.E.M. *Significant decrease from the paired control day, p < .05. Duloxetine had less effect than gabapentin on the responses in the control group*.

### Analgesic responses in sham group

In general, the effects of the different analgesic drugs were similar in the GEL^TM^ and sham groups whenever the sham group demonstrated pain behavior; five of the 8 rats in the sham group eventually developed pain behaviors like the GEL group. Morphine demonstrated marked analgesia to the pinprick stimuli on the ipsilateral side on day 28, but not on later days. This correlates with the waning effect of morphine in the GEL group. Celecoxib did not affect responses in the sham group. As in the GEL group, both gabapentin and duloxetine demonstrated marked analgesia in the sham group.

### Summary of analgesic effects

Morphine depicted a waning analgesic response over time not related to tolerance, with increasing pain behaviors, suggesting of a developing opioid-induced hyperalgesia. Celecoxib had no analgesic effect on pain behaviors at any time. By contrast, gabapentin and duloxetine both produced robust analgesia bilaterally in the GEL group during all time periods and in 5/8 shams during the last two drug testing periods. The pain behaviors, when present in the GEL group after D23 and later in the shams after D90, respond to all analgesics similarly.

## Reversal of GEL effect with EPO

The pain behavior in the GEL group that had persisted for 4 months was reversed for up to 7 days, (end pilot) by the targeted application of epoetin alfa in a pilot study, at the end of the investigation. After the main experiment was over on day 149, the 14 GEL^TM^ animals continued in a pilot study lasting 8 days. Three subgroups were formed: 1.)” “at-site” EPO group, n=5, 2.) subcutaneous EPO injection group, n=4, and 3.) control group n=5. Under isoflurane, as described prior, the “at-site” group received injections of epoetin alfa (EPO) 200 units in normal saline (NS) at the site of the original GEL procedure. The subcutaneous group received a subcutaneous EPO injection of 200 units in NS (n=4), and the control group received no injection. Days 140 and 149 after the GEL procedure were taken as reference days and then EPO was injected as a perineural infiltration once in the EPO groups on day 152 only on the left leg. Two of the rats in the” “at-site” EPO group received a second local EPO injection with a lateral approach on day 155 (seen as a+sign on the left paw results Fig 12). Pinprick behavior data were collected on days 152, 153, 154, 156, 159, and 160. The resulting data were analyzed using a mixed model ANOVA, with the EPO injection group as the between subjects factor (3 groups), and days as one repeated measures factor (8 days), and laterality as a second repeated measures factor (left and right paws). The data are presented in Fig 12). The three-way interaction was not significant, but the groups-by-days interaction was significant (*F* (14, 77)=8.208, *p*<.001). As can be seen in the graph, the effect of EPO was nearly identical on the right and left sides and the effect was significant for at least 6 days, end of study. An ANOVA such as this should be interpreted with caution because of several cells with zero variance (there is a ceiling effect of 8 paw withdrawals out of 8 stimulus presentations of pinprick).

**Fig 12.**
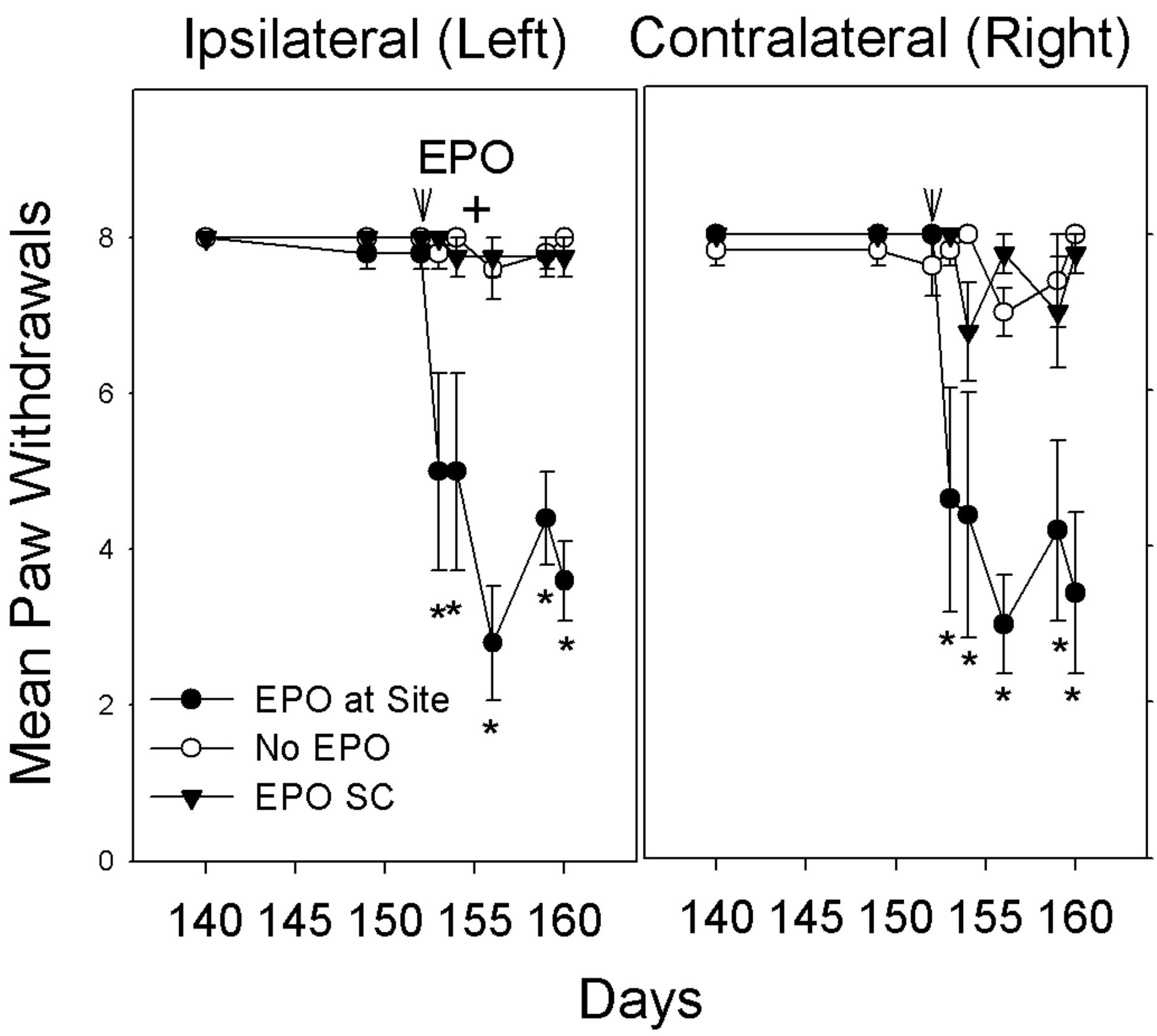
Pain behavior decreased bilaterally after EPO was injected at the GEL site (n=5) on day 152 only on the left leg. *After a perineural infiltration of EPO (200 units in NS) the paw withdrawals to pinprick (hyperalgesia or hypersensitivity) decreased bilaterally to near pre-GEL procedure levels. The groups of “No EPO"” (n=5) and “EPO SC” injected subcutaneously (n =4) continued with robust pain behavior, bilaterally. The+on day 155 signifies two GEL rats still with pain behavior that had the GEL site on left re-injected with EPO with a different anatomic technique, as described, which caused a decrease in their paw withdrawals to pinprick. *p <.001 EPO at Site Group vs. No EPO*.

## Results of histology

The tissue sections from 9 rats randomly selected from each procedure group were blinded, and observed by an independent neuropathologist. These observations were later respectively matched to each of the three groups, with n=3 GEL, n=3 sham, and n=3 the controls. Details are described in Histology within the Methods section. In brief, there were no differences noted in the tissue sections between the control and the sham-treated animals, and no abnormal findings in either group. The structure of the nerve and surrounding tissue was completely unremarkable (Fig 13).

**Fig 13.**
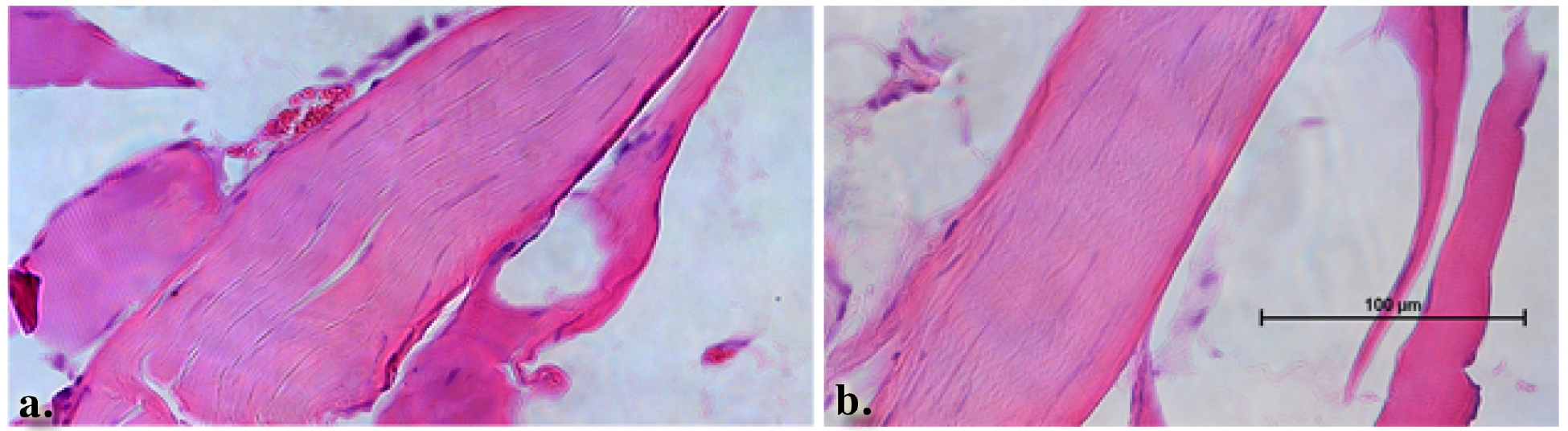
Normal tibial nerve histology in both the control and sham procedure groups. *Longitudinal sections through the control (a.) and sham (b.) procedure groups revealed normal appearing nerves. The surrounding muscle tissue that was harvested with the nerve also appears to be within normal limits*.

Within the GEL^TM^ treated group, the histology of the nerves was in stark contrast to either of the control groups. The gross appearance of the nerves at the time of dissection revealed a discrete area of swelling, or a bulge, along the course of the distal tibial nerve, in all specimens of the GEL group only. These discrete structures were about twice the diameter of the nerve just distal and proximal to the outcropping. Longitudinal sections through the portion of the tibial nerve that contained these bulges revealed that the swelling is the result of changes within the endoneurium, including evidence of intraneural edema with increased spacing between the neural fascicles, and axonal edema in the fibers within the bulge region (Fig 14a). and Fig 14b). In addition, there were numerous profiles in which ongoing axonal fragmentation, a hallmark of Wallerian degeneration, was evident (see arrows in Fig 14a) and Fig 14c) (66).

**Fig 14.**
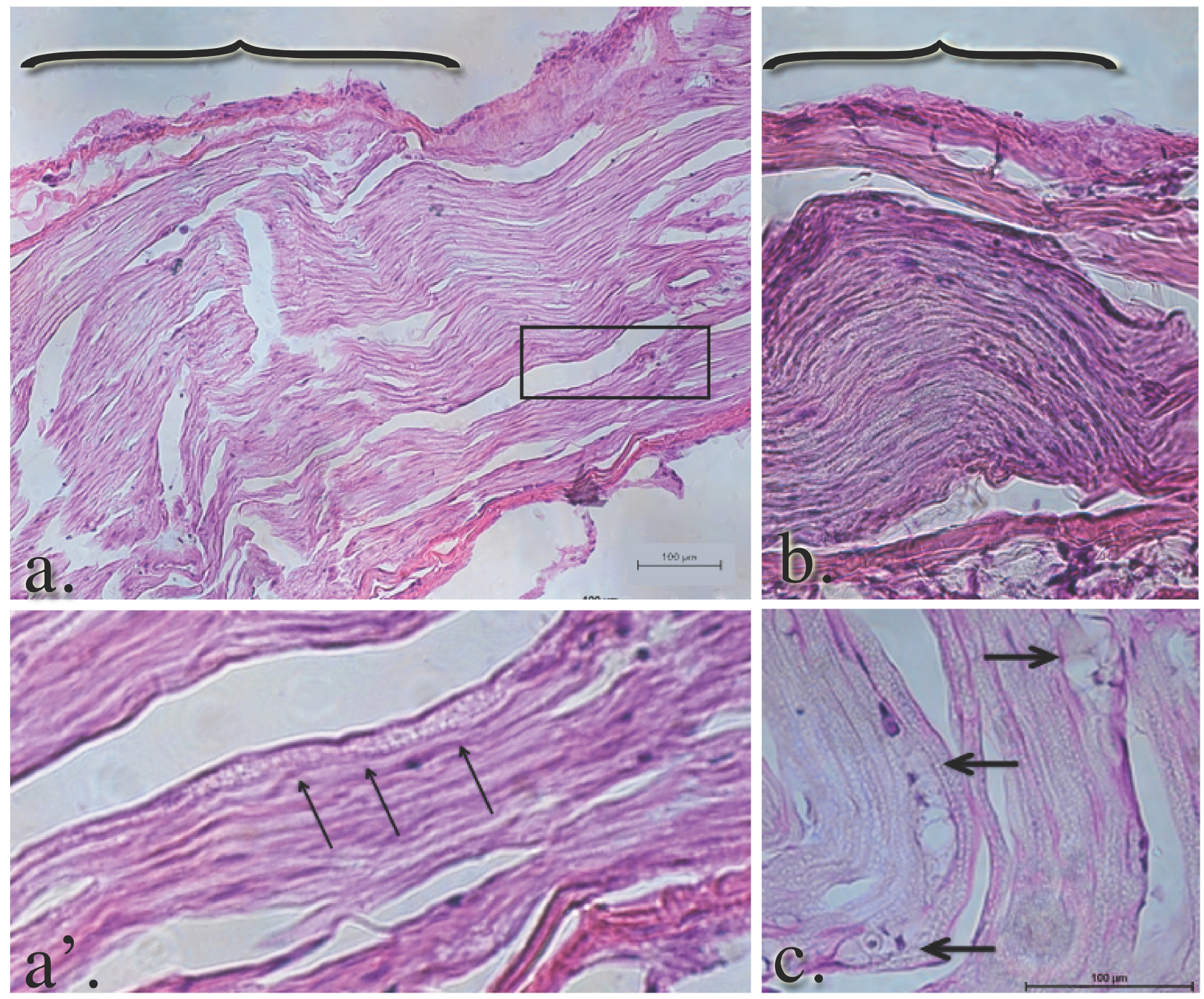
Edema and Wallerian degeneration in the nerves of GEL procedure rats. *Panels a. and b. reflect the gross swelling seen in the tibial nerves upon dissection. The brackets in panels a. and b. denote the areas of swelling in the GEL nerves. The axons within the swollen area are themselves swollen. The diameters of the axons in this distended area of the nerve are approximately twice the diameter of the contiguous axons proximal or distal (not shown) to it. The arrowheads point to axonal debris in panel a which is a magnification of the area within the box in panel a. Macrophage-engulfled myelin and axonal debris (panel c, arrows) is further evidence of ongoing Wallerian degeneration.*

Consistent with ongoing Wallerian degeneration, we observed numerous macrophages within the endoneurium, which phagocytize myelin and axonal debris (Fig 15a). and Fig 15b). large arrows), we also noted a significant leukocyte accumulation in and around the perineurium (Fig 15b). thin arrows).

**Fig 15.**
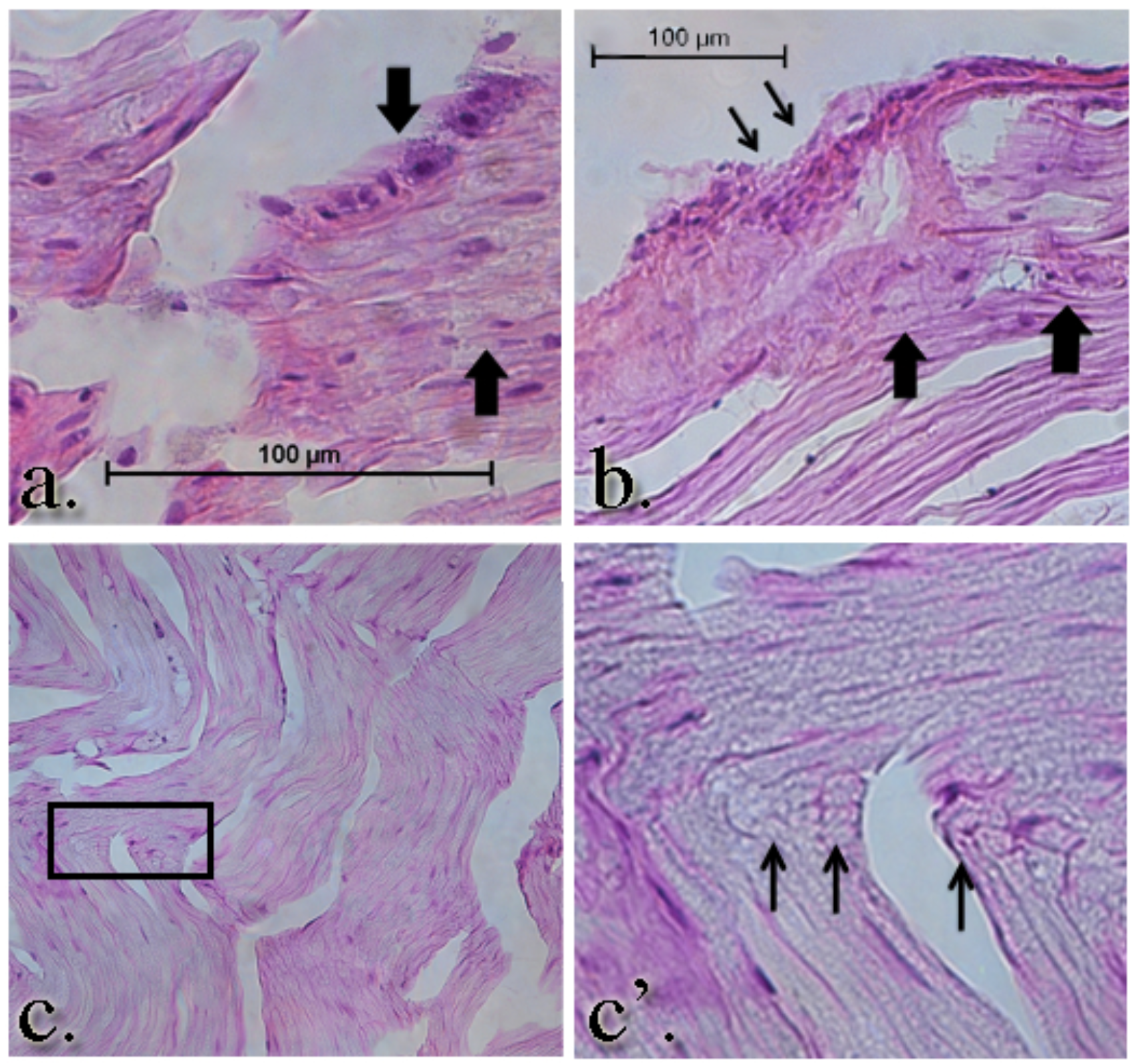
Simultaneous Wallerian degeneration and regeneration is evident in the GEL group nerves. Intraneural (large arrows a. and b.) and perineural leukocytic (small arrows, b.) infiltration is seen in the GEL group nerves. Notably, there are numerous profiles of regeneration axons in the same nerve (panel c. and c') at the same proximal/distal level, suggesting ongoing nerve remodeling in the GEL procedure animals. Panel c’ is a higher power magnification of the clusters of small, unmyelinated axons within the box in panel c.

Findings consistent with any gel residual were not noted in the GEL specimens. In addition to the ongoing Wallerian degeneration observed in the GEL™ animals, we also noted ongoing axonal regeneration. There were clusters of small, unmyelinated or lightly myelinated fibers growing into the nerves of the GEL cohort only. The grouping of these small fibers is consistent with regeneration, rather than the randomly single-arrayed, unmyelinated fibers that are found in the healthy, homeostatic adult nerve (67).

### A factor influencing pain behavior testing

A reportable factor discovered during this study, relates to the influence of hormone replacement of the experimenter on the elicited response to stimuli. On each routine pain behavior test day, after the first 6 random rats were screened, the results were compared to the prior session of each rat, to observe for environmental influences. In nine such early screening comparisons, paw withdrawals were markedly less to nonexistent bilaterally in the 6 rats prescreened as compared to their prior screening session. Thirty minutes after the topical application to the investigator of a 17 beta-estradiol replacement cream, the screening was repeated on the same 6 rats in each screening, and their elicited pain behaviors were then consistent with the data collected during the prior testing period. On unblinding, this effect was not related to groups. Even baseline pre-procedure behaviors were similarly affected by the estrogen hormone replacement. All data used in this study was collected with topical estrogen applied 30 minutes prior to beginning all testing. This finding needs further study.

## Discussion

The nonsurgical NeuroDigm GEL™ model appears to mimic many humans complaining of persistent neural pain, by being of gradual onset, persisting for months and lacking deformities or antalgic gait. Two other characteristics have been suggested for relevance in a rodent neuropathic pain model. These are an analgesic response profile similar to humans, and effective analgesia with doses of near human equivalent strengths (S4 a). This socially mature rodent chronic pain model closely approaches these analgesic response guidelines.

While tissue injury causes“nociceptive pain” or acute pain, the tissue repair or tissue healing that follows may produce neuropathic pain. Chronic pain has been defined as starting after 3-6 months in humans or after the healing of a tissue injury (68). This temporal delay correlates to the last phase of tissue repair, called tissue remodeling (69), occurring months after the initial inflammatory stages of tissue repair following an acute soft tissue injury (45), (70), (71). By inducing tissue remodeling, this model accelerates the usual delayed onset of neuropathic pain. The pain behavior in this model is robust at day 23, rather than beginning over months as in humans (S1). Analgesic screening can be started on day 23.

We used older rats in this study to look at age as a factor in the manifestation of chronic neural pain (72–76). Currently, most neuropathic pain studies use rats weighing 200 to 300 grams with ages of 50 – 67 days or a human equivalent of 13 – 14.5 years, an average age conversion being in the adolescence period (59), (77), allowing for sexual maturity. The rats in this study started at 9.5 months of age and ended at 15.4 months of age, which calculates to a 24-year-old to a 39-year-old human equivalent using the average age rate conversion formula for the adult phase (59). The mature adult rats were chosen to resemble more closely, human chronic pain patients (1), (78).

Performing analgesic studies on a mature adult rat with a physiologic neural lesion may reveal more accurately the actual analgesic potential of an agent, and therefore confirm the reliability of the rodent as a representative animal model for neuropathic pain.

Central sensitivity or neural plasticity is demonstrated in this model after 2-3 weeks by the appearance of elicited pain behaviors on the contralateral side in all GEL^TM^ rats, as well as the 5/8 sham rats that developed the late onset pain behavior. Central sensitivity with contralateral pain is known to exist in humans with originally unilateral neural pain (79–82; however, contralateral pain is not commonly documented in most rodent models of traumatic nerve lesions.

The behavioral data suggests that opioid-induced hyperalgesia (83–86) with increasing neural pain behavior appears to develop over time, as seen in the GEL and sham groups (Fig 8). Such lack of effective analgesia with morphine is characteristic of many neuropathic pain patients (87), (88). Since each of the three doses of morphine in this study is separated by weeks, the weakening response to morphine does not reflect tolerance. Preclinical repeated screenings of analgesics (89) over several weeks may be more relevant for translational effectiveness (90). This model may have negative predictive validity for morphine. Morphine's decreasing analgesia may correlate to the neural injury (93) or developing neural regeneration. Gabapentin and duloxetine demonstrated analgesia after one dose in each testing period for the GEL and sham groups. Clinical experience has shown that analgesia with one dose is common with duloxetine and gabapentin in many adults, especially the elderly (S4 b).

Our present study with socially mature male rats (greater than 6 months old (59) revealed the hyperalgesia as the more robust and persistent of the pain behaviors, as compared to the allodynia in this aged model. Our NeuroDigm GEL™ method was reproduced prior in younger rats (270g), the mechanical allodynia appeared robust, and much higher doses of gabapentin and duloxetine were used (S4 c). Many elderly patients (>70 years old), with chronic nerve pain, have been found to have mechanical hyperalgesia more commonly than mechanical allodynia (S4 b). More studies are needed to discern how age affects pain behaviors (72) and the effective dose of an analgesic.

Erythropoietin (91–94), methylprednisolone (95), glucocorticoids in general (96), and ARA290, an erythropoietin derived tissue repair peptide (97), (98), as well as other biologics (99), are known to have neuroprotective effects systemically and locally in nerve injury rodent models. A surgical proximal sciatic nerve defect model in a prior study showed the myelin repair potential of a sustained localized erythropoietin delivery method (100). The local erythropoietin dose at the neural lesion, in this model may integrate the tissue repair ability of erythropoietin (101) in alleviating pain behaviors and regaining normal neural function.

The targeted local injection of an erythropoietin analog in this study appears to support the localized“ectopic” and “generator” theories for the persistence of neural pain, as the unilateral application of erythropoietin appears to have eliminated, for at least a week (end of study), the bilateral pain behavior of mechanical hyperalgesia (pinprick). The focal neural swelling seen on the distal tibial nerve in the GEL group may act like a mid-axon nociceptive stimulus, maintaining the pain behavior until the localized erythropoietin healed (S4 d) the neural lesion.

An unanticipated feature suggested in our five-month study is the evidence for habituation to the von Frey and brush stimuli. In this study, the light touch stimuli (von Frey, brush) have less painful significance than the pinprick, so their responses may be susceptible to habituation. We have not been able to locate any specific study of habituation to von Frey or brush stimuli. Responses in the GEL group appeared to diminish in post-procedure periods 4 and 5, with the difference between the GEL and control groups maintained at a significant level. The effect is seen clearest in the summary variable in on brush stimuli in Fig 4). By contrast, the sham group tended to remain the same or increase during the same time periods, as some of them developed pain behavior. We did not observe any evidence of habituation to the pinprick stimulus in any group.

The nerve specimens used in this study were processed at end of the 5 month study, where chronic tissue changes dominated, without acute inflammation (102). The histologic changes seen were restricted to the NeuroDigm GEL™ procedure group and are consistent with ongoing tissue remodeling in the area where the GEL was placed. Specifically, there was evidence of both ongoing Wallerian degeneration and axonal regeneration, hallmarks of nerve remodeling. Neural remodeling can be a response to compression (103). Also consistent with ongoing nerve remodeling is the observed inflammation with leukocytic infiltration, seen in both the endoneurial environment and the extraneural space.

Notably, a large number of tightly packed unmyelinated fibers were within the nerves of the GEL procedure animals, consistent with regeneration. The increased number of these fibers and their unusual clustered appearance raises the issues of adequate insulation and the possibility that some of the pain behavior might be due to ephaptic transmission (104–106). Neural regeneration with ephaptic transmission is likely the underlying cause of the both the Tinel's sign (107) observed in some patients at sites of neural compression due to entrapment, and also the Pinch Reflex Test found at sites of regenerating peripheral nerves in experimental rodents (108), (109).

The histology demonstrates that the light microscopy anatomical response to the GEL procedure is restricted to a focal area of neural and axonal edema with neuroinflammation in the tibial nerve, yet the behavioral effects are widespread. The contralateral spread of pain behaviors in the GEL model raises the possibility that there are central changes as well. Whether this is the result of anatomic changes or the alterations in neural signaling remains to be determined. Occasionally, as seen in our shams and the paclitaxel model (110), lesions visible by light microscopy need not be present to manifest neuropathic pain behavior. Additional studies are needed in which the effects of the GEL^TM^ distal peripheral nerve procedure on the thalamus, the DRG, and spinal cord are explored.

The mechanisms that lead to the anatomic and behavioral changes are not yet determined. However, it is likely that the neural remodeling (96) seen in our model reflects many of the ongoing changes that are seen with chronic pain following peripheral trauma (111). The findings in our study are consistent with prior findings found in neural biopsies of humans with severe persistent pain due to known nerve entrapments (107), (112), (113). The creation of this model allows for further in-depth studies to understand the events around the establishment of chronic neural pain following minimal trauma as seen in humans.

An unexpected and powerful influence of topical estrogen on the investigator was noted in this study. The postmenopausal status of the investigator appeared to markedly diminish the paw withdrawal response in all groups. However, this effect was reversed within 30 minutes by the investigator applying a 17 beta-estradiol topical cream. This reaction echoes the olfactory ‘male observer’ effect of male experimenters reducing acute pain behaviors in rodents, as compared to females (114). This hormonal finding suggests that besides their sex, the age and hormonal status of experimenters may influence the reproducibility of pain behaviors.

## Conclusion

Our model was created to simulate a focal sustained compression (a pinch) on a nerve, which has long been considered a cause of persistent neural pain (115), (116). Gradual perineural changes of the extracellular matrix and scarring (117) can cause such focal compressions on a nerve (40).

Recently, a lesion of the somatosensory system, or a disease, has been needed for the definition of neuropathic pain (118). This model provides an extended temporal window into a neural lesion as a naturally occurring soft tissue disease.

We have shown how a percutaneous procedure can induce neuropathic pain with a delayed onset as usually found in humans. This refined method supports the 3Rs initiative for the humane use of animals, and is clinically relevant. A biomimetic mononeuritis can be induced by stimulating the physiology of tissue repair (119), (120). This tissue reaction after trauma is an encoded orchestrated response in the peripheral nervous system, as in all vertebrate soft tissues. The last stage of this repair process is tissue remodeling, (121), (122) which may be a unifying etiology (89) in many deceptively complex neuropathic pain syndromes.

Crucially, our model depicts opioid-induced hyperalgesia as seen in humans—without causing surgical injury to animals. We have used tissue repair in this model to create neuropathic pain, and also to heal it. The refined (123) NeuroDigm GEL™ model with a target can help discover biomarkers, effective analgesics, lesion detection devices, targeted biologic treatments and alternatives to opiates.

S1 Supplement 1: Tissue repair comparison graph

S2 Supplement 2: Dataset

S3 Supplement 3: Detailed Anova statistics on groups

S4 Supplement 4: Other References

S5 Supplement 5: ARRIVE guidelines

## Acknowledgements

We are indebted to general surgeon Dale C. Rank Sr. M.D. (deceased February 5th 1996), who contributed to the translation from man to the preclinical method of the model. Deep gratitude is given to Gordon Munro PhD, who advised on analgesic doses and methods, and read the paper.

## Author Contributions

Conceived and designed the experiments: MRH DAF DEW JLB. Performed the experiments: MRH RMD DEW. Analyzed the data: DAF DEW MRH. Wrote the paper: MRH DAF RMD DEW JLB

